# Functional, Immunogenetic, and Structural Convergence in Influenza Immunity between Humans and Macaques

**DOI:** 10.1101/2025.02.21.639368

**Authors:** Maya Sangesland, Ning Li, Yaroslav Tsybovsky, Megan D. Rodgers, Julianna Han, Alesandra J. Rodriguez, James A. Ferguson, Amy R. Henry, Sarah C. Smith, Jesmine Roberts-Torres, Rebecca A. Gillespie, Cuiping Liu, Jonah S. Merriam, Tyler Stephens, Connor Williams, Emma Maestle, Martin Corcoran, Michelle Ravichandran, Adrian Creanga, Sarah F. Andrews, Theodore C. Pierson, Gunilla B. Karlsson Hedestam, Chaim A. Schramm, Douglas S. Reed, Daniel C. Douek, Tongqing Zhou, Andrew B. Ward, Masaru Kanekiyo

## Abstract

Human B cell immunity to the influenza hemagglutinin (HA) stem region, a universal influenza vaccine target, is often stereotyped and immunogenetically restricted, posing challenges for study outside humans. Here, we show that macaques vaccinated with a HA stem immunogen elicit human-like public B cell lineages targeting two major conserved sites of vulnerability, the central stem and anchor epitopes. Central stem antibodies were predominantly derived from V_H_1-138, the macaque homolog of human V_H_1-69, a V_H_-gene preferentially used in human central stem broadly neutralizing antibodies (bnAbs). Similarly, macaques produced anchor bnAbs with the human-like NWP motif. Both bnAb lineages were functionally and structurally analogous to their human counterparts, with recognition mediated largely by germline-encoded motifs. Thus the macaque immunoglobulin repertoire supports human-like public bnAb responses to influenza HA. Moreover, this underscores the utility of homologous germline-encoded immunity, suggesting that immune repertoires of macaques and humans may have been similarly shaped during evolution.

**HIGHLIGHTS:**

● Functional human-like public antibody lineages can be elicited to HA stem supersites in macaques.
● Macaque central stem bnAbs are predominantly derived from V_H_1-138, a V_H_-gene homologous to human V_H_1-69.
● The human-like CDR L3 NWP anchor epitope-targeting lineage can be elicited in macaques.
● Central stem and anchor bnAbs from humans and macaques engage their respective epitopes with atomic level similarity.

## INTRODUCTION

The immunoglobulin repertoire contains exceptional diversity, where affinity for a given antigen is mediated by the B cell receptor (BCR). Each BCR contains both V-gene encoded and hypervariable complementarity determining regions (CDRs), the latter which are formed by the rearrangement of V(D)J gene segments during B cell development^1^. Of the CDR loops that compose the antibody paratope, the third heavy chain CDR (CDR H3) contains the majority of BCR diversity and consequently serves as the primary determinant of antigen specificity^2^. Despite this diversity, there are recurrent examples of stereotyped or public antibody responses whereby antigen recognition is mediated by germline-encoded motifs, including V-gene encoded CDRs, allowing for the reproducible elicitation of structurally identical antibodies across genetically unrelated individuals^3^. Moreover, these public responses can center upon highly conserved and broadly protective epitopes of hypervariable viruses such as HIV-1, influenza, or SARS-CoV-2, providing genetic templates for vaccine design^4–6^.

Influenza virus remains a global public health concern, posing an ongoing challenge due to continuous antigenic evolution and pandemic potential^7^. The primary vaccine antigen and target of humoral immunity is the surface glycoprotein hemagglutinin (HA), which divides influenza A viruses (IAVs) into two antigenically and phylogenetically distinct groups (group 1 and group 2). Structurally, HA is composed of a hypervariable head domain and a less variable stem region; the latter contains several functionally conserved epitopes targeted by heterosubtypic broadly neutralizing antibody (bnAb) lineages^8^. However, the HA stem is immunologically subdominant, with stem-directed B cell responses consistently maintained at sub-protective levels, both following infection or vaccination^9–11^. This has prompted efforts to direct the humoral response to conserved features through rationally designed HA stem-only immunogens as well as chimeric HA constructs^12,13^.

While subdominant, humans can generate public bnAb lineages to the HA stem with broad activity within or across IAV HA groups. A large proportion of central stem BCRs use the human V_H_1-69 gene, where the unique V_H_-gene encoded CDR H2 loop contains affinity to the hydrophobic groove within the central stem epitope^14–17^. This germline-encoded affinity has been shown to provide a reproducible source of epitope specificity which can then be readily activated and amplified through vaccination with the correct germline-stimulating immunogen^11,18^. The HA stem also contains another antigenic supersite, termed the anchor epitope, which is membrane proximal^19–21^. In humans, anchor antibodies are highly stereotyped, requiring germline-encoded motifs to engage the target epitope. Importantly, both central stem and anchor bnAb lineages were elicited in humans upon seasonal influenza vaccination, followed by either a chimeric HA (cHA) or H1 stabilized stem ferritin nanoparticle (H1ssF) vaccine; NCT03300050 and NCT03814720, respectively^22,23^, suggesting their clinical relevance in anti-influenza immunity.

Nonhuman primates, particularly macaques, are widely used in vaccine studies due to their evolutionary proximity to humans^24^. While macaques have been shown to generate functional serological immunity following vaccination with HA stem-based vaccines^25–27^, there is currently minimal evidence that human-like pathways for bnAb elicitation is possible. In general, public B cell immunity has not been well characterized outside of humans, thus understanding if macaques are genetically hardwired for such responses not only provides translational value for vaccine evaluation but also insight into the evolutionary utility of germline-encoded immunity.

In this study, we investigated the B cell response following vaccination with H1ssF nanoparticle in cynomolgus macaques. We found that macaques can recapitulate human HA stem immunity, generating public antibody lineages with specificity to the central stem and anchor epitopes. Isolation and single-cell sequencing of BCRs unveiled that central stem antibodies were overwhelmingly derived from the macaque V_H_1-138 gene, the closest homolog to human V_H_1-69. Anchor antibodies were more diverse but did contain the canonical human-like lineage possessing the CDR L3 NWP motif. Antibodies from both lineages were broadly neutralizing, protective, with paratopes consisting of germline encoded and structurally convergent motifs. Our data thus indicate that B cell immunity in macaques can recapitulate human public bnAb responses to the HA stem at the functional, immunogenetic, and structural levels. More broadly, the shared immunoglobulin gene usage in response to two distinct epitopes suggests that the immunoglobulin repertoires of humans and macaques may have been shaped similarly during evolution.

## RESULTS

### Induction of central stem and anchor epitope-specific serum neutralizing antibodies

We utilized historic samples from cynomolgus macaques (*Macaca fascicularis*) immunized with H1ssF, a nanoparticle vaccine displaying stabilized HA stem-only trimers, adjuvanted with AF03^25^ (Figure 1A). In vaccinated animals, the titer of HA stem-specific serum IgG increased upon each immunization with H1ssF, which occurred concomitant with increased microneutralization titer and breadth across IAV reporter viruses, including vaccine-matched A/New Caledonia/20/1999 (H1N1 NC99), unmatched but within subtype A/Michigan/45/2015 (H1N1 MI15), and heterosubtypic H5N1 A/Vietnam/1203/2004 (H5N1 VN04) and H2N2 A/Singapore/1/1957 (H2N2 SI57) (Figures 1B-C).

**Figure 1.**
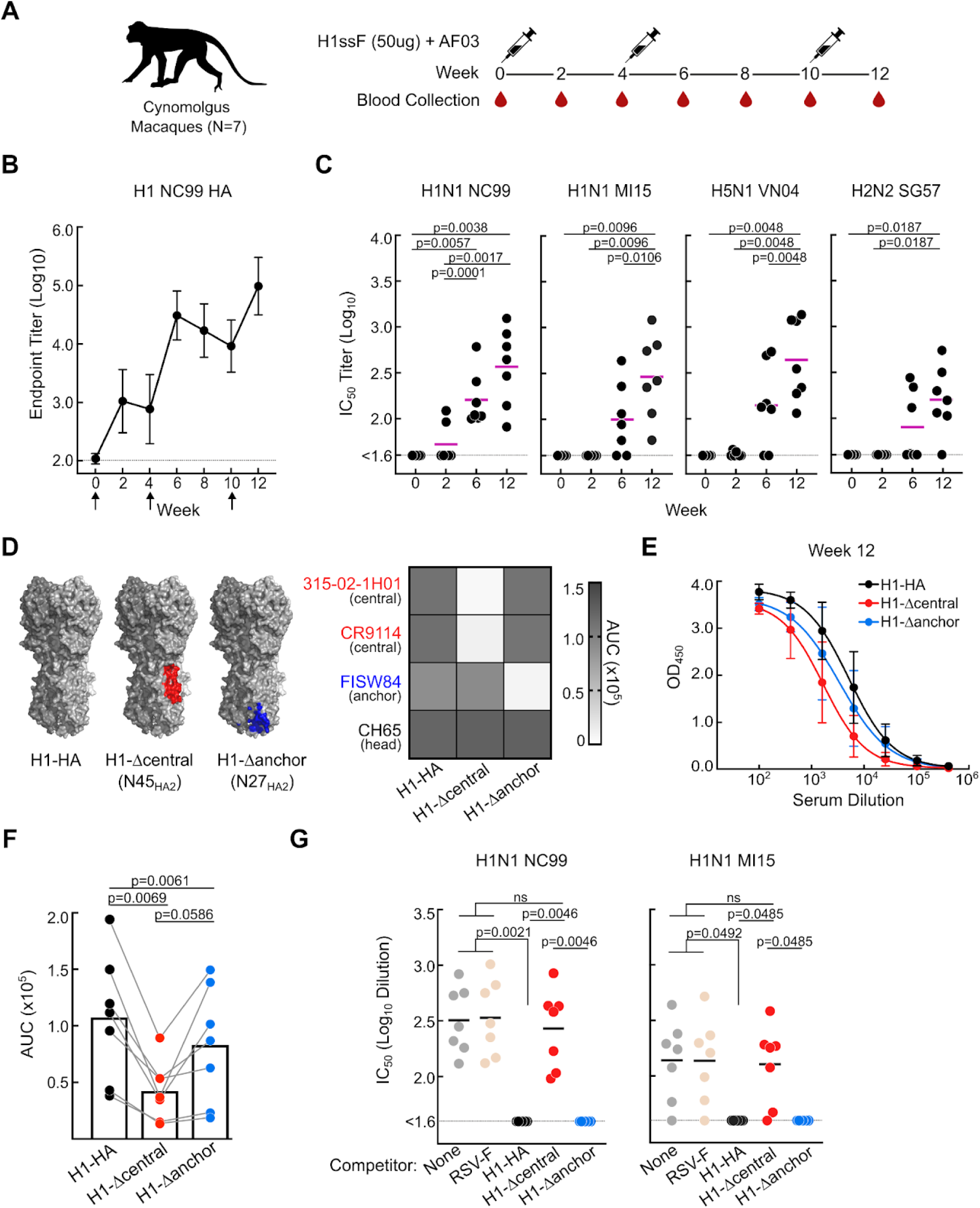
Serum epitope specificity and neutralizing activity in macaques following H1ssF immunization. **(A)** Sequential immunization scheme with H1ssF. Influenza naive cynomolgus macaques (n = 7) were vaccinated thrice with H1ssF and AF03 adjuvant^25^. **(B)** Serum IgG endpoint titer to recombinant H1 (NC99) HA trimeric protein at wks 0, 2, 4, 6, 8, 10 and 12. Arrows denote vaccination timepoints. Horizontal line denotes limit of detection. N = 7 animals, mean and SD are shown. **(C)** Serum neutralizing antibody response against H1N1 A/New Caledonia/20/1999 (NC99), H1N1 A/Michigan/45/2015 (MI15), H5N1 A/Vietnam/1203/2004 (VN04), and H2N2 A/Singapore/1/1957 (SG57) reporter viruses^60^. Purple line denotes mean. Statistics performed by one-way ANOVA with Geisser-Greenhouse correction and Tukeys post hoc test. **(D)** Representation of H1 NC99 HA trimer probes: WT HA containing central stem and anchor epitopes (H1-HA), addition of a N45_HA2_ glycan (red, H1-Δcentral), and addition of a N27_HA2_ glycan (blue, H1-Δanchor). Central stem (red, PDB: 3GBM) and anchor (blue, PDB: 6HJQ) epitopes are mapped on HA (PDB: 3LZG) (Top). Binding (AUC) of representative head (CH65), central stem (CR9114 and 315-02-1H01), and anchor (FISW84) mAbs against H1-HA, H1-Δcentral, H1-Δanchor probes (Bottom). Samples run in duplicate. **(E-F)** Serum IgG response at week 12 to H1-HA (black line), H1-Δcentral (red line), and H1-Δanchor (blue line). N=7 animals, mean and SD are shown (E). Corresponding serum binding (AUC) at week 12. Statistical significance determined by one-way ANOVA with Geisser-Greenhouse correction and Tukeys post hoc test (F). **(G)** Microneutralization (IC_50_) titer of immune sera at week 12 against H1N1 NC99 and H1N1 MI15 pre-absorbed with the following competitors: none (grey), RSV-F (tan), H1-HA (black), H1-Δcentral (red), H1-Δanchor (blue). Statistical significance determined by one-way ANOVA with Geisser-Greenhouse correction and Tukeys post hoc test (n = 7 animals, line denotes mean).

To assess if macaques can elicit polyclonal responses to the conserved central stem and anchor epitopes, we applied a series of HA trimer probes, including the full-length trimeric ectodomain of H1 NC99 (H1-HA), H1 NC99 containing a glycan at residue N45_HA2_ (H1-Δcentral) or a glycan at position N27_HA2_ (H1-Δanchor) to block access to the central stem^28^ or anchor epitopes^21^, respectively (Figure 1D). We found that polyclonal serum antibodies to both epitopes were present in vaccinated macaques as evidenced by the reduction in binding to the H1-Δcentral and H1-Δanchor probes as compared to H1-HA (Figures 1E-1F and S1A-B). Serum antibodies to the central stem were more prevalent than to the anchor epitope at each post-vaccination timepoints at weeks 2, 6, and 12 (Figures 1E-F and S1A-B), and contributed to almost all detected serum neutralizing activity against vaccine-matched H1N1 NC99 and unmatched MI15 reporter viruses (Figures 1G and S1C). While not reaching the 50% cutoff, polyclonal anchor antibodies did show weak neutralizing activity against the NC99 reporter virus (Figure S1C). Thus, macaques can generate functional polyclonal serum antibodies to both the central stem and anchor epitopes.

### Immunogenetic characteristics of macaque HA stem B cell lineages

To evaluate the H1ssF-induced B cell response, we analyzed peripheral blood mononuclear cells (PBMCs) by flow cytometry from four macaques (BB798E, 6974, T009 and R996) after the final H1ssF immunization (week 12). Total HA specific B cells, as determined by staining with a H1-full length trimer probe (H1-FL), were readily detected (1.7%) relative to IgG (Figures 2A and S1D). However, upon staining with H1-stem and H1-full length trimer probes (H1-stem^+^/H1-FL^+^), herein referred to as HA^+^, the antigen specific B cells segregated into distinct upper and lower clusters (Figures 2B, S1D).

**Figure 2.**
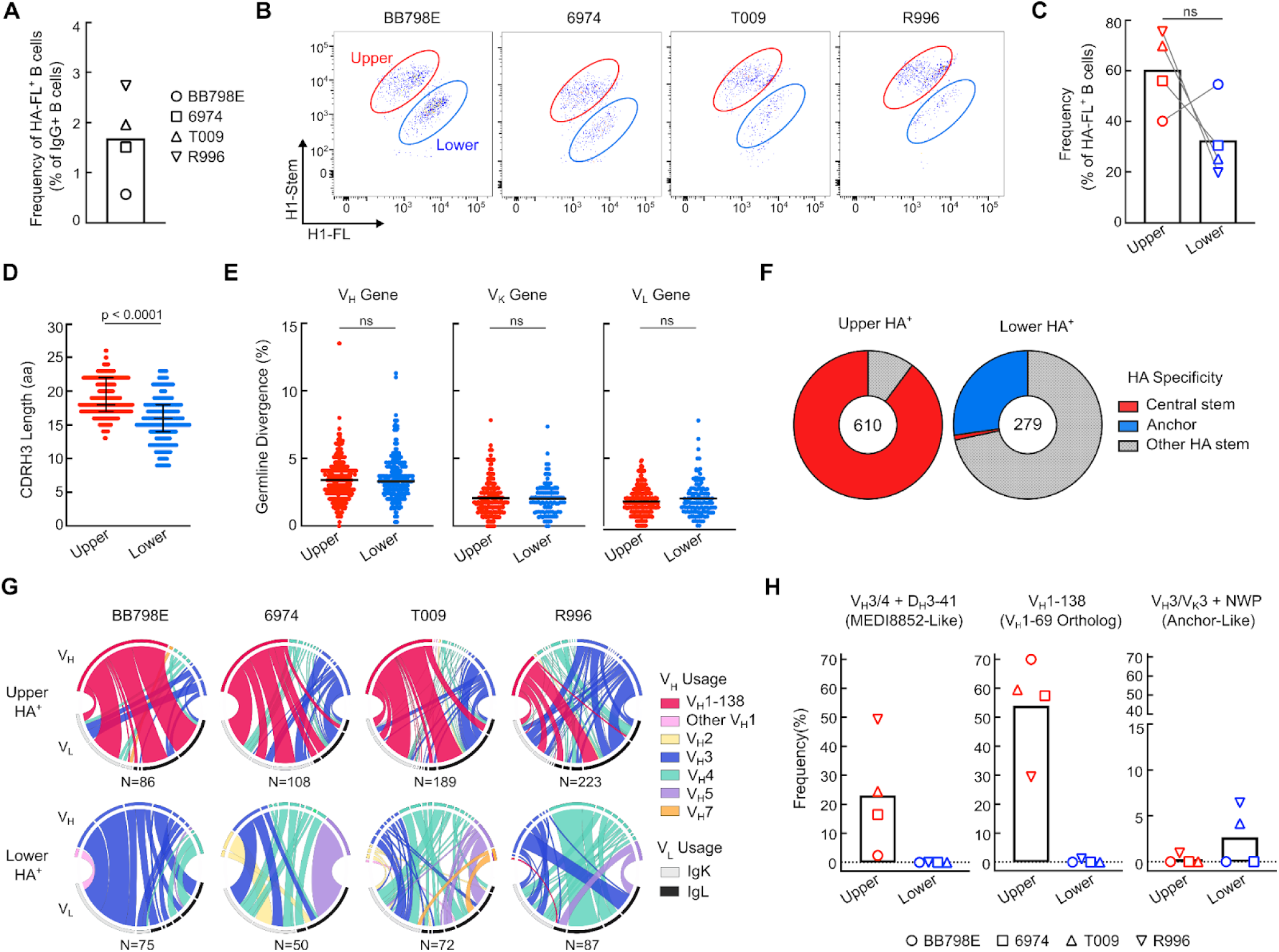
Antigenic and immunogenetic characteristics of HA specific B cells. **(A)** Frequency of antigen specific B cells reactive to H1-FL at week 12, relative to IgG^+^ B cells. H1-FL^+^ B cells are defined as CD3^-^/CD14^-^/CD16^-^/CD56^-^/CD20^+^/IgG^+^/IgM^-^/H1-FL^+^ (see Figure S1D for gating). Symbols denote individual animals. **(B-C)** Flow plots of HA^+^ (H1-stem^+^/H1-FL^+^) upper cluster (red) and lower cluster (blue) B cells across 4 macaques (B). Corresponding frequency of each upper and lower cluster as a percent of H1-FL reactive B cells (see also Figure S1D for gating). N = 4 animals are shown, symbols denote individual animals. Statistical significance determined by Wilcoxon matched-pairs signed rank test (C). **(D)** CDR H3 length distribution for upper and lower HA^+^ clusters. Shown are median and interquartile range. Significance determined by two-tailed Mann-Whitney test. **(E)** Percent germline divergence across V_H_, V_K_ and V_L_ chains in each corresponding upper and lower cluster. Significance determined by two-tailed Mann-Whitney test. **(F)** Antigenicity of upper and lower HA^+^ sorted single B cells from RATP-Ig supernatants. Antigenicity was determined by binding to H1-HA, H1-Δcentral, and H1-Δanchor probes (see also Figure 1D for probes). Antibody supernatants that did not bind H1-Δcentral or H1-Δanchor probes were denoted as “Other HA stem”. **(G)** Chord diagrams showing V_H_ (top half) and V_L_ (bottom half) gene pairs in HA^+^ B cells from the upper and lower clusters. Total number of sequence pairs are listed. **(H)** Frequency of the relevant lineages; MEDI8852-Like: V_H_3/4+D_H_3-41, V_H_1-69 homolog: V_H_1-138, and human anchor-like: V_H_3/V_K_3+NWP, across 4 macaques (symbols denote individual animals).

Both clusters contributed to the majority of HA-FL reactive B cells (∼62% upper vs. ∼35% lower), with the upper cluster in higher frequency in 3 out of 4 macaques (Figures 2B-C and S1D). Macaques also generated H1-stem single positive B cells, suggesting the presence of another immunogenic surface on the stem-only antigen, such as the trimer apex formed by the removal of the HA head (Figures S1E-F).

To assess the antigenic and immunogenetic characteristics of the upper and lower HA^+^ clusters, we isolated HA-specific B cells and applied the Rapid Assembly, Transfection, and Production of Immunoglobulins (RATP-Ig) workflow to screen antigen specificity through recombinant monoclonal antibody (mAb) expression in parallel with BCR sequencing^29^. While CDR H3 lengths differed between the BCRs in the upper and lower clusters (upper: 13-26 aa, lower: 9-23 aa), the overall percent germline divergence (somatic hypermutation, SHM) was unchanged within the heavy and light chain genes of each cluster (Figures 2D-E). However, striking differences were observed in the antigen specificities, where ∼90% of B cells in the upper cluster targeted the central stem epitope compared to ∼1% of B cells in the lower cluster (Figure 2F). Anchor targeting B cells were present exclusively in the lower cluster at ∼30%, a population dominated by non-central/non-anchor ‘other HA-stem’ specificities (Figure 2F).

Within the upper HA^+^ cluster, we observed an overwhelming bias for B cell clones containing the macaque V_H_1-138 V_H_-gene (KIMDB nomenclature, http://kimdb.gkhlab.se)^30^, which was present across all four vaccinated animals, encompassing 29–70% of the total V_H_-genes within that cluster (Figures 2G-H). V_H_1-138 is the closest macaque homolog to human V_H_1-69, a V_H_-gene most commonly utilized by human central stem-targeting bnAb lineages^31^. While prevalent in humans, macaques were previously thought unable to generate the V_H_1-69 bnAb lineage as they lacked the critical hydrophobic I53/F54 CDR H2 residues uniquely encoded by the V_H_1-69 V_H_-gene that is required for HA stem recognition in humans^14^. Within the central stem we also identified the previously characterized macaque bnAb lineage containing a V_H_3 or V_H_4 family heavy chain gene paired with various D_H_3-41*01 alleles (herein referred to as V_H_3/4+D_H_3-41), which shares resemblance to the human V_H_6-1+D_H_3-3 class of bnAbs including MEDI8852^25,26^ (Figures 2G-H).

The composition of the lower cluster was more heterogeneous but did contain BCRs with a V_H_3 heavy chain paired with a V_K_3 light chain along with a CDR L3 NWP motif (hereby referred to as V_H_3/V_K_3+NWP); characteristics associated with human anchor epitope-targeting antibodies^20,21^. V_H_3/V_K_3+NWP BCRs were present in two out of four animals, albeit at low frequencies (Figure 2H). These results suggest that macaques may employ human-like public responses against two distinct bnAb epitopes on the HA stem following immunization with H1ssF.

### Functionality of macaque central stem and anchor antibody lineages

We next assessed the functionality of the B cell response by expressing vaccine-elicited central stem and anchor BCRs as recombinant mAbs. Akin to humans, macaque central stem mAbs neutralized multiple influenza reporter viruses within group 1 IAVs, including homotypic vaccine matched H1N1 NC99 and heterologous H1N1 MI15, H5N1 VN04, and H2N2 SG57 viruses (Figure 3A). In contrast, anchor mAbs were less broad overall, a feature shared with their human counterparts^20,21^, neutralizing only H1N1 viruses (Figure 3A). We did not detect any neutralization against group 2 HA viruses, consistent with a prototypic group 1 response elicited by H1ssF^23^ (Figure 3A). Of the central stem mAbs, only non-V_H_1-138 mAbs neutralized the H2N2 reporter virus. Human H2N2 HAs contain a bulky F45_HA2_ residue within the stem domain, instead of an I45_HA2_, which typically occludes access of V_H_1-69-class antibodies to this site^32,33^, and we predict this may contribute to loss of macaque V_H_1-138 reactivity against this virus. Moreover, the vaccine-elicited mAbs retained neutralizing activity at potencies comparable to prototypic human central stem and anchor bnAbs (Figure 3B), suggesting that eliciting functional antibodies to these epitopes in macaques is not dependent upon repeated virus exposure, prior seasonal influenza vaccination, or in the case of the anchor epitope, the membrane anchored display of HA.

**Figure 3.**
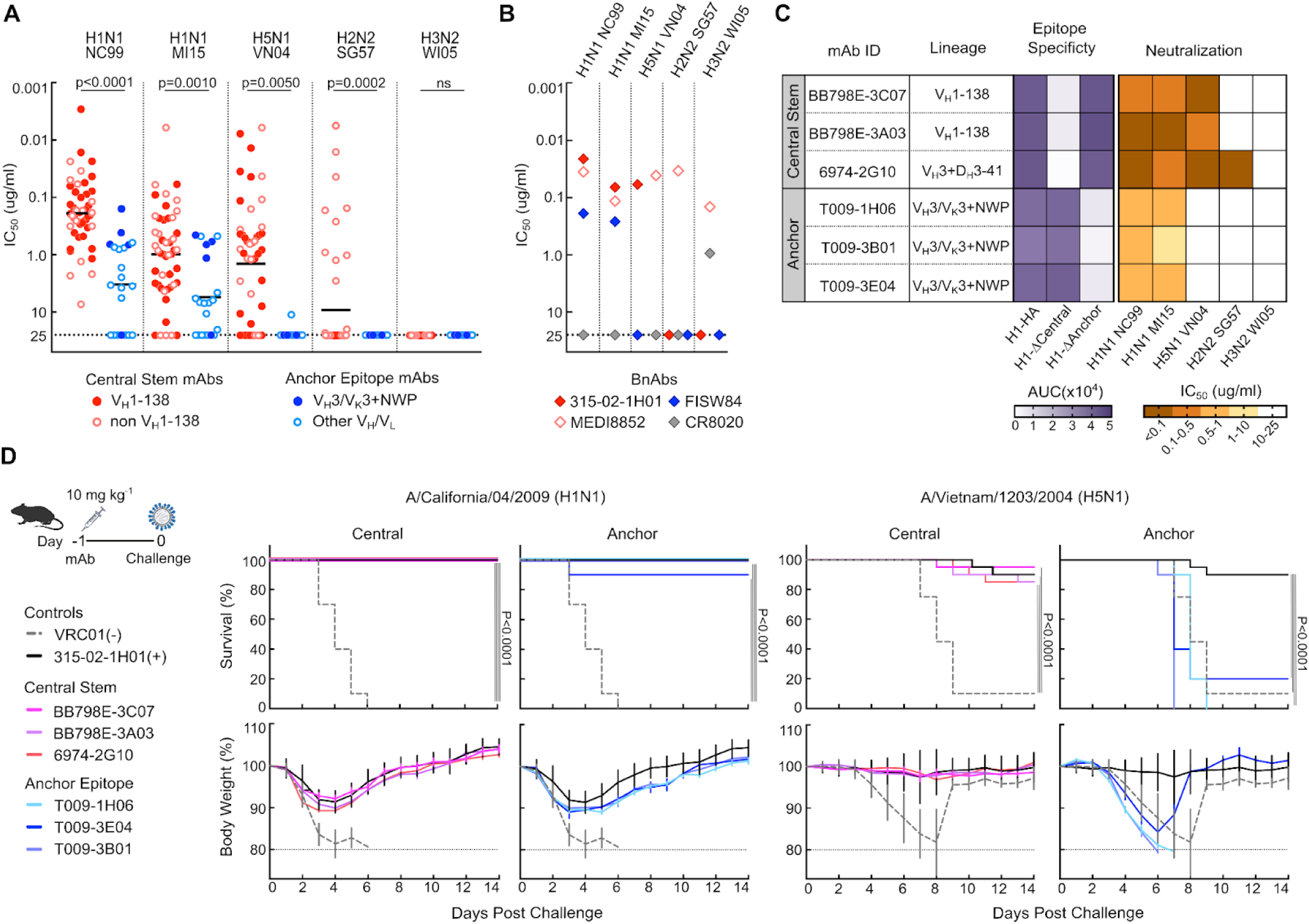
Macaque central stem and anchor antibodies share functional characteristics to their human counterparts. **(A)** Microneutralization (IC_50_) activity for macaque H1ssF elicited monoclonal antibodies against H1N1 A/New Caledonia/20/1999 (NC99), H1N1 A/Michigan/45/2015 (MI15), H5N1 A/Vietnam/1203/2004 (VN04), H2N2 A/Singapore/1/1957 (SG57), and H3N2 A/Wisconsin/67/2005 (WI05) reporter viruses. Central stem (red) and anchor (blue) mAbs are denoted. Bar corresponds to mean. Statistical significance determined by two-tailed Mann-Whitney test. **(B)** IC_50_ values of prototypic influenza stem bnAbs. Central stem: 315-02-1H01 (human V_H_1-69 bnAb) and MEDI8852; Anchor: FISW84; Group 2 stem: CR8020. **(C)** Characteristics of 3 central stem and 3 anchor macaque bnAbs. Lineage, epitope specificity, and microneutralization titer are shown (see also Figure S2A). **(D)** Pre-exposure prophylaxis experiment with central stem (BB798E-3C07, BB798E-3A03, and 6974-2G10) and anchor (T009-1H06, T009-3B01, T009-3E04) antibodies. BALB/c mice were treated with 10 mg kg^-1^ of mAbs via the intraperitoneal route 24 h prior to intranasal virus challenge with H1N1 A/California/04/2009 and H5N1 A/Vietnam/1203/2004 viruses. Percent survival and percent body weight change are shown. Error bars indicate SD. Multiple comparisons of Kaplan-Meier curves were performed by the log-rank test with Bonferroni correction.

To investigate protective efficacy, we selected three central stem (BB798E-3C07, BB798E-3A03 and 6974-2G10) and three anchor mAbs (T009-1H06, T009-3E04, and T009-3B01), encompassing the relevant macaque HA stem immunoglobulin lineages, for *in vivo* protection studies (Figures 3C and S2A). When given prophylactically, all mAbs provided protection against vaccine-unmatched H1N1 A/California/4/2009 infection in mice (Figure 3D). However, only central stem mAbs protected against heterosubtypic H5N1 VN04 challenge, consistent with our *in vitro* neutralization profiles (Figures 3A and 3C-D). Collectively, these findings confirm that in addition to convergent immunogenetic characteristics, macaque central stem and anchor antibodies are functionally analogous to human HA stem bnAb lineages.

A feature of some influenza HA stem antibodies is their tendency for polyreactivity^34–36^. We thus assessed the polyreactivity of macaque central stem and anchor mAbs by binding to cardiolipin and HEp-2 cells. While control mAbs behaved as expected^37^, we found little to no reactivity of the macaque mAbs to these antigens (Figures S2B-C).

### V_H_1-138 bnAbs engage the HA central stem through germline-encoded structural motifs

We next compared the sequence characteristics of macaque V_H_1-138 central stem bnAbs with previously identified human V_H_1-69 central stem bnAbs^33^, where bnAbs as defined as those that, at minimum, neutralize both H1N1 and H5N1 reporter viruses. Human V_H_1-69 antibodies are characterized by a hydrophobic CDR H2 which encodes an apex I53/F54 motif used to engage the hydrophobic groove in the epicenter of the central stem^14,15,38^. While known macaque V_H_1-138 alleles do not encode for the canonical human I53/F54 motif, the mAbs and bnAbs derived from them do contain a hydrophobic CDR H2 apex with either a germline-encoded L53/V54 or a somatically mutated L53/A54 (Figures 4A and S3A). Macaque V_H_1-138 bnAbs also exhibited centralized CDR H3 tyrosine residues, a hallmark of the human V_H_1-69 bnAb lineage^38^(Figure 4A). CDR H3 tyrosine residues were present across V_H_1-138 central stem BCRs as a whole but not within other central stem macaque lineages (Figure S3B). Additionally, the SHM of macaque V_H_1-138 central stem bnAbs averaged ∼3.7%, which was lower than the ∼5.8% SHM found in the previously identified human V_H_1-69 central stem bnAbs^33^, likely due to successive rounds of affinity maturation from prior influenza exposure in humans (Figure 4B).

**Figure 4.**
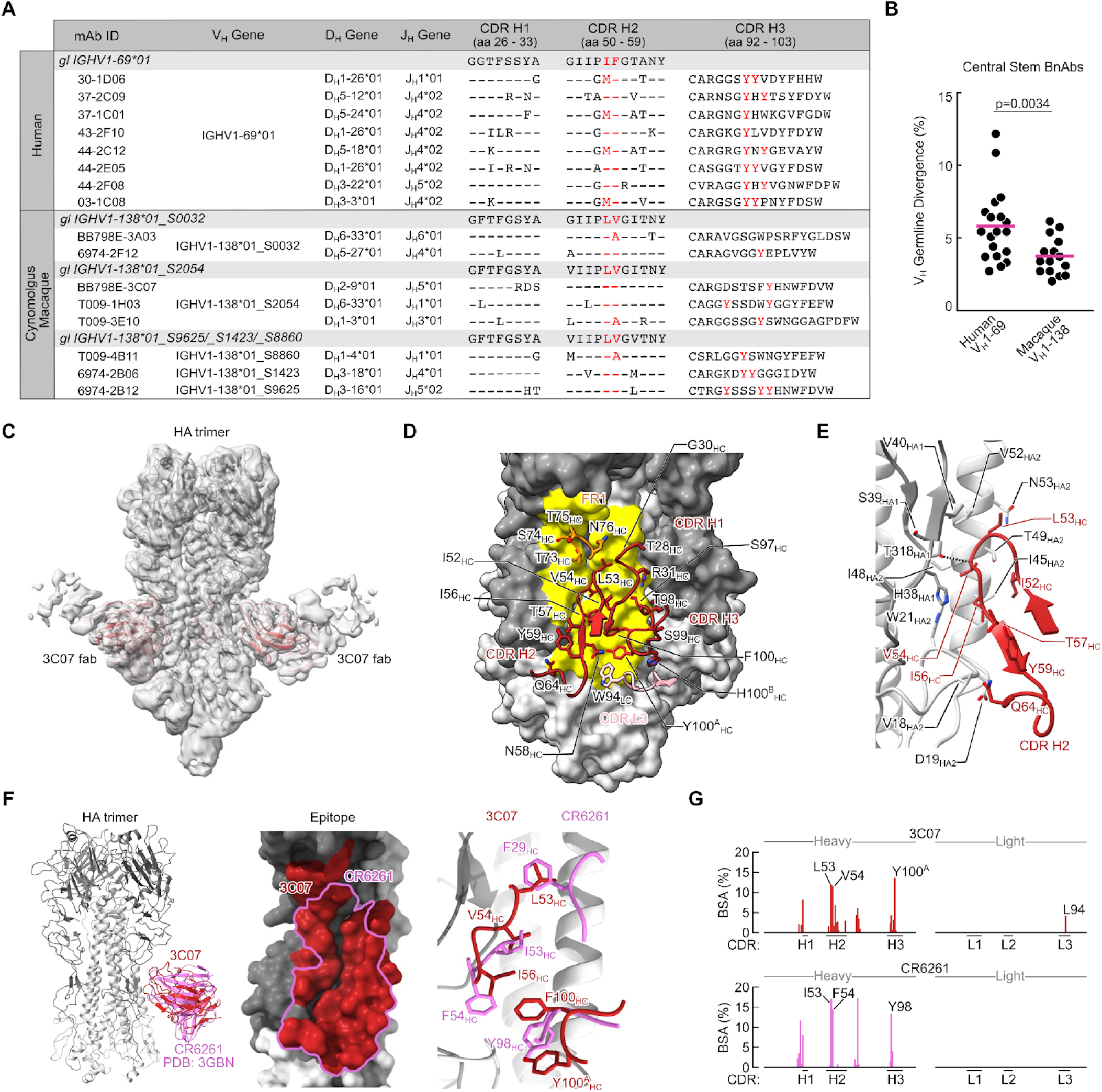
IGHV1-138 central stem bnAb lineages and structural analysis. **(A)** Heavy chain genetic characteristics of human IGHV1-69*01^33^ and macaque IGHV1-138 bnAbs. Central stem bnAbs are defined as those that neutralize H1N1 and H5N1 reporter viruses. The corresponding germline alleles and sequences for CDR H1 and CDR H2 are shown as reference. Critical CDR H2 residues at position 53/54 and CDR H3 tyrosine residues are highlighted in red. Kabat numbering is indicated for CDR H1, CDR H2, and CDR H3. **(B)** Percent germline divergence (nucleotide level) of previously identified human V_H_1-69 bnAbs^33^ compared to macaque V_H_1-138 bnAbs. P values were calculated using a two-tailed Mann-Whitney test. **(C)** Cryo-EM map of BB798E-3C07 in complex with H1 NC99 HA trimer. Heavy and light chains of the Fab are colored in red (HC) and light pink (LC). **(D)** Contact interface between HA (surface representation) and BB798E-3C07. CDR regions of Fab and residues in contact with HA are labeled. The contact area on HA is colored yellow. **(E)** Detailed illustration of Fab BB798E-3C07 CDR H2 interaction with HA. Only molecular regions participating are shown for clarity. Dashed lines depict hydrogen bonds. **(F)** Superposition of Fab BB798E-3C07 with the crystal structure of Fab CR6261 (PDB: 3GBM) (Left). Comparison of the contact interface between BB798E-3C07 and CR6261 (Middle). Superposition of residues of BB7998E-3C07 Fab and CR6261 Fab forming key contacts with HA (Right). BB798E-3C07 is shown in red and CR6261 is shown in orchid. **(G)** Contributions of each CDR to total BSA for BB798E-3C07 and CR6261. Only CDRs participating in the interaction are shown.

To investigate the molecular basis for public V_H_1-138 usage in macaque central stem antibodies, we applied cryogenic electron microscopy (cryo-EM) to determine the structure of the V_H_1-138 bnAb BB798E-3C07. Cryo-EM structure of BB798E-3C07 Fab in complex with H1 NC99 HA trimer was solved at 3.7 Å resolution (Figures 4C-E, S4A-G, and Table S1). We found that BB798E-3C07 engages the central stem with a slightly downward angle predominantly using its heavy chain, akin to HA stem recognition exhibited by human V_H_1-69 bnAbs^15,16^. CDR H2 contributes the majority of the antibody paratope with germline-encoded residues L53_HC_, V54_HC_ and I56_HC_ mediating contact to the hydrophobic groove on the HA stem (Figure 4E). Specifically, L53_HC_ interacts with helix A residues V52_HA2_ and N53_HA2_ as well as HA1 residue V40_HA1_. Residue V54_HC_ interacts with W21_HA2_ residing at the epicenter of the hydrophobic groove as well as I48_HA2_ and T49_HA2_ on helix A. I56_HC_ also interacts with W21_HA2_ and I45_HA2_. CDR H3 Y100^A^_HC_ interacts with I45_HA2_ as well as G20_HA2_, T41_HA2_ and Q38_HA2_ (Figures 4D-E). The only light chain interaction occurs through CDR L3 W94_LC_ to D19_HA2_ (Figure 4D). Lastly, germline reverted gHgL [germline-H (gH) and germline-L (gL)] BB798E-3C07 Fab maintained binding to H1 NC99 HA, further highlighting germline affinity to the HA stem (Figure S3C-D).

We next compared the antibody paratopes of BB798E-3C07 and CR6261, a prototypic human V_H_1-69 central stem bnAb^16^. Both bnAbs were heavy chain centric in their mode of recognition, sharing a similar footprint burying a total surface area of 962 and 813 Å^2^ on HA, respectively (Figure 4F).

Importantly, we found that the key CDR H2 contacts were maintained across both bnAbs where the I53_HC_ and F54_HC_ interactions of CR6261 were replaced by V54_HC_ and I56_HC_ in BB798E-3C07. In addition, interactions made by F29_HC_ and Y98_HC_ in CR6261 were replaced by L53_HC_ and F100_HC_/Y100^A^ in BB798E-3C07 (Figure 4F). The per residue buried surface area (BSA) contributions were also remarkably similar between the two bnAbs with CDR H2 making more than one third of the contributions for both BB798E-3C07 and CR6261 (Figure 4G). Overall, these results indicate that akin to human V_H_1-69 bnAbs, macaque V_H_1-138 bnAbs can accommodate contacts mediated by germline-encoded residues, resulting in highly convergent recognition to the central stem.

### V_H_1-69 homologs are found across the primate order

V_H_1-69 homologs are widely present across the primate order with the highest degree of homology to V_H_1-69*01, an allele frequently used in human central stem bnAbs^17^, within great apes, old- and new-world monkeys (Figure 5A). Such widespread homology suggests that these V_H_-genes likely arose from a common primate ancestor. While it is unknown if other primates are able to elicit a ‘V_H_1-69-like’ germline-encoded response to the HA stem, we find that the hydrophobic CDR H2 is maintained in V_H_1-69 homologs amongst great apes, old-, and new-world monkeys, a defining feature shared with human V_H_1-69 stem bnAbs (Figures 5A and S5A). To assess if the CDR H2 domains of other non-human primates can accommodate binding to HA, we replaced the CDR H2 domain of germline reverted gHgL BB798E-3C07 with the CDR H2 regions of select V_H_1-69 primate homologs. We find that binding to HA NC99 is largely maintained in these ‘swap’ antibodies when the corresponding CDR H2 residues at position 53 and 54 were hydrophobic, and where position 54 was not a phenylalanine, a key encoded feature of human V_H_1-69 bnAbs (Figure 5B-C). Additionally, we found that the presence of a hydrophobic residue at position 56 aided in the overall binding stability to HA (Figures 5B-C). While CDR H2 F54 is critical for human V_H_1-69 recognition of the HA stem, we predict that F54_HC_ within macaque BB798E-3C07 will clash with I48_HA2_ and T49_HA2_ on HA helix A, contributing to the loss of binding of the human CDR H2 swap gHgL BB798E-3C07 to the central stem (Figures 5B-C and S5B).

**Figure 5.**
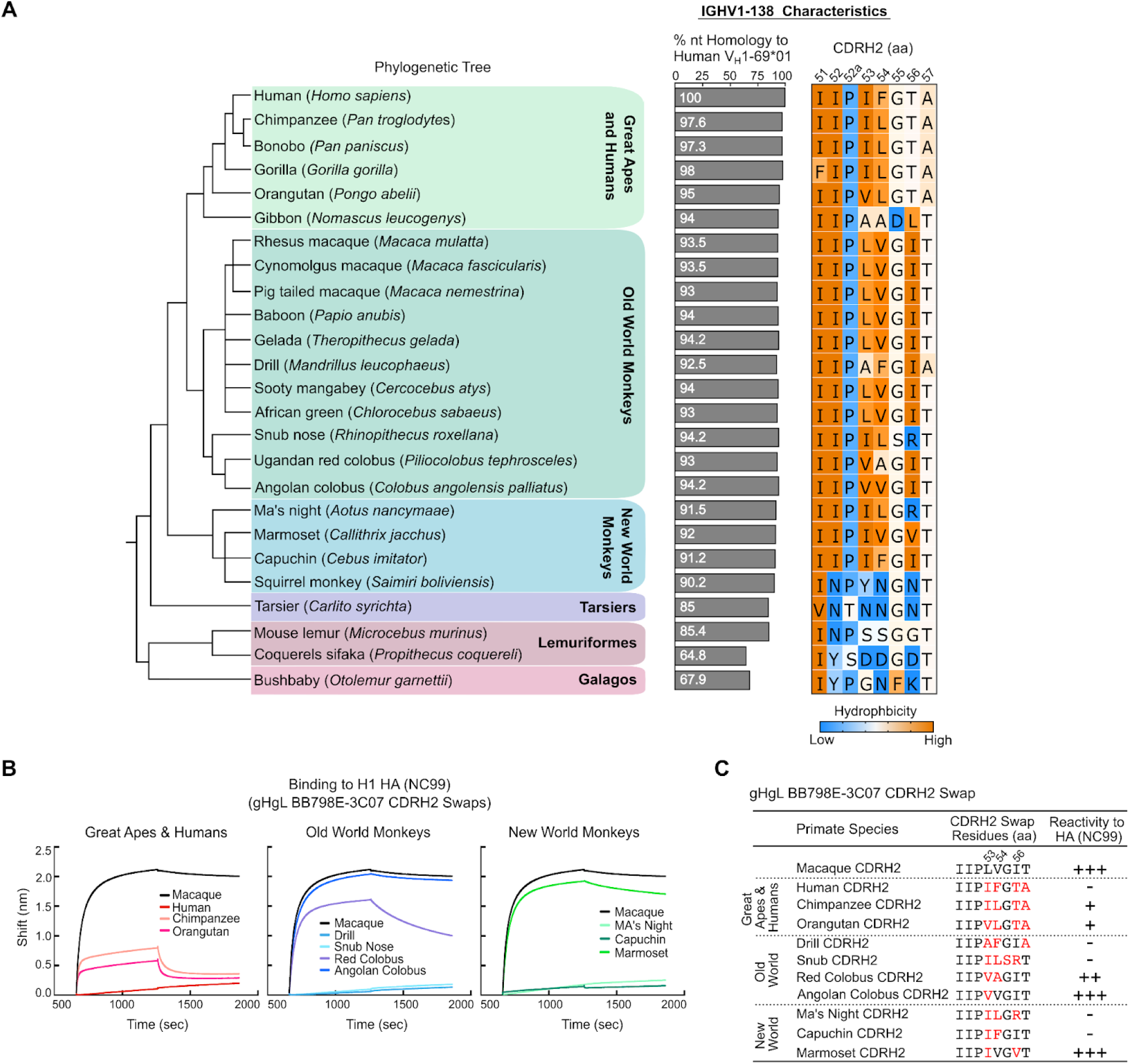
Human IGHV1-69 homologs are found across the primate order. **(A)** Phylogenetic relationship of primate species based on the NCBI taxonomy browser (Left). Percent nucleotide sequence homology of primate IGHV1-69*01 homologs compared to the human IGHV1-69*01 V_H_-gene (Middle). Corresponding CDR H2 amino acid residues within each primate IGHV1-69 homolog colored by hydrophobicity (orange: high, blue: low) based on the scale of Kyte and Doolittle^84^ (Right) . **(B)** BLI sensogram of gHgL BB798E-3C07 ‘CDR H2-swap’ mAbs binding to H1 NC99 HA. Swap mAbs are derived from gHgL BB798E-3C07 where the original macaque CDR H2 domain of BB798E-3C07 is replaced by the CDR H2 sequence from a corresponding V_H_1-69 primate homolog. All binding curves are compared to macaque gHgL BB798E-3C07. **(C)** Table comparing the gHgL BB798E-3C07 swap mAbs and their corresponding CDR H2 sequences (see also Figure 5A for CDRH2 sequences). Red denotes residues changed from the original macaque CDR H2 sequence. +/- denotes strength of binding to H1 NC99 HA.

### Macaque V_H_3/V_K_3+NWP antibodies share genetic and structural characteristics of human anchor bnAbs

Human anchor antibodies are highly stereotyped and characterized by use of a V_H_3 heavy chain gene (V_H_3-23, V_H_3-48 or V_H_3-30) paired with a V_K_3 light chain gene (V_K_3-15 or V_K_3-11) and a germline-encoded CDR L3 NWP motif^20,21^. While low frequency, we do identify anchor antibodies in two H1ssF vaccinated macaques sharing these features (Figure 2G). At the sequence level, these antibodies use homologous V_H_, V_L_, and J_L_ gene segments and maintain all critical residues defined in human anchor bnAbs, namely the CDR H2 Y58_HC_ and NWP motif within CDR L3 (Figure 6A). Additionally, we identified a functional anchor antibody which uses a CDR L3 SWP motif replacing the canonical NWP (Figure 6A).

**Figure 6.**
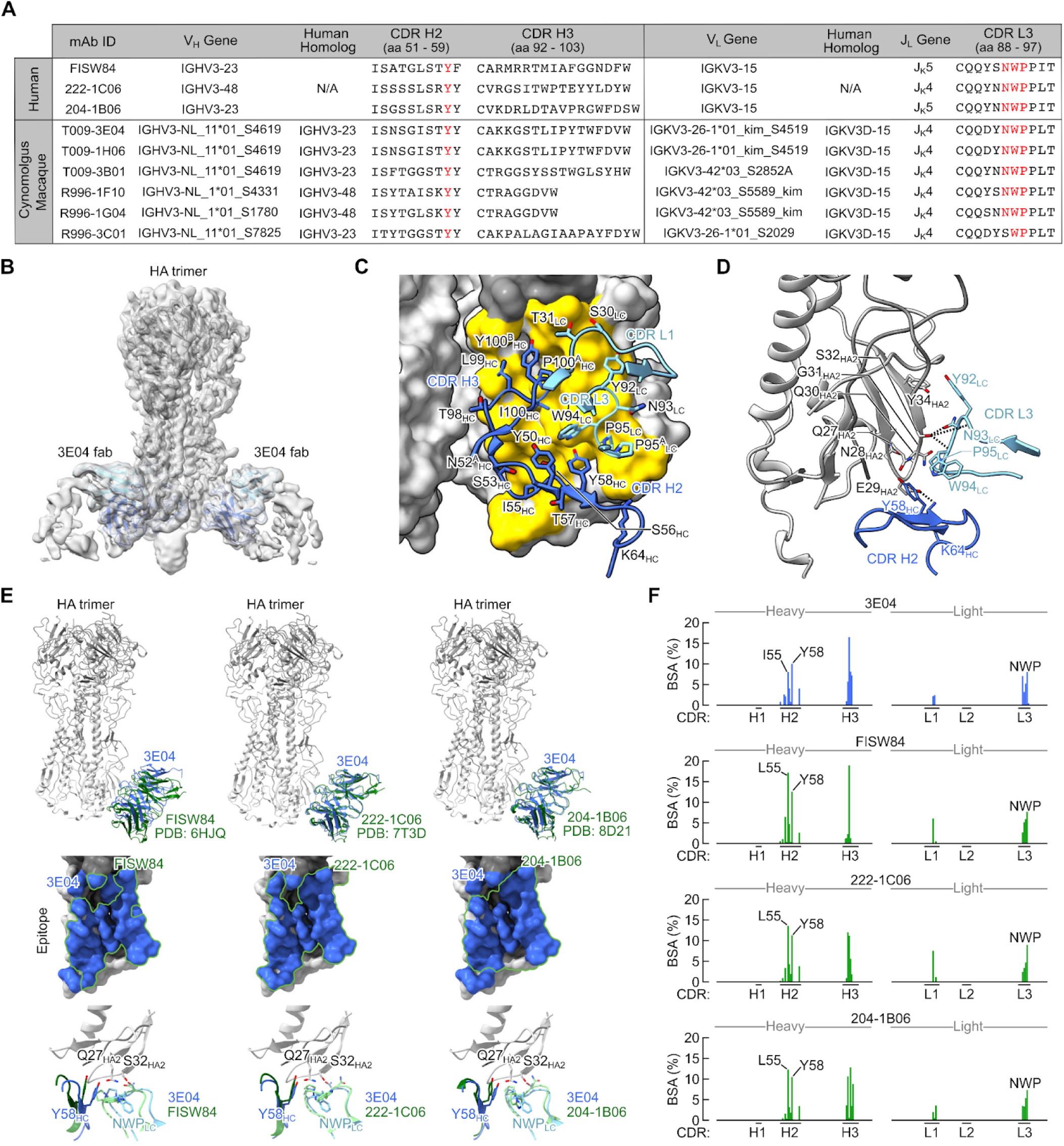
Structural characteristics of the anchor bnAb T009-3E04 in complex with HA. **(A)** Heavy and light chain genetic characteristics of human and macaque anchor antibodies. Critical residues are highlighted in red. Kabat numbering is indicated. **(B)** Cryo-EM map of Fab T009-3E04 in complex with H1 NC99 HA trimer. Heavy and light chains of Fab are colored in blue (HC) and light blue (LC). **(C)** Contact interface between HA (surface representation) and T009-3E04. CDR regions of Fab and residues in contact with HA are labeled. The contact area on HA is colored yellow. **(D)** Detailed illustration of HA and T009-3E04 with participating interactions shown. Dashed lines depict hydrogen bonds and salt bridges. Colors are consistent with panels B and C. **(E)** Superposition of T009-3E04 with prototypic human anchor antibodies FISW84 (PDB: 6HJQ), 222-1C06 (PDB: 7T3D) and 206-1B06 (PDB: 8D21). Comparison of the contact interface between T009-3E04 (colored in blue) and FISW84, 222-1C06, and 206-1B06 (outlined in green) is shown. **(F)** CDR Contribution to total BSA for T009-3E04, FISW84, 222-1C06, and 204-1B06. Only participating CDRs are shown.

To characterize the mode of HA recognition of a macaque V_H_3/V_K_3+NWP anchor mAb, we determined the cryo-EM structure of T009-3E04 Fab in complex with H1 NC99 at 3.8 Å (Figures 6B-D, S6A-G, and Table S1). T009-3E04 binds HA near the membrane proximal region with a slightly upward angle (Figure 6B). The paratope of T009-3E04 was composed primarily of the CDR L3 NWP motif and the CDR H2 and H3 loops (Figures 6C-D), with a total BSA of 834 Å at the Fab–HA interface. While CDR H3 mainly engages residues within HA1, CDR H2 and L3 loops make contacts with HA2 near the base centered around S32_HA2_ and Q27_HA2_ (Figure 6D). There are hydrogen bonds between the hydroxyl group of S32_HA2_ and three CDR L3 residues (Y92_LC_, N93_LC_, and W94_LC_) as well as between E29_HA2_ and CDR H2 Y58_HC_ (Figure 6D). We also performed negative stain EM (nsEM) on R996-3C01 Fab, an anchor antibody containing a non-canonical CDR L3 ‘SWP’ motif, and we find that despite not having the ‘NWP’, this Fab engaged a similar epitope. Indeed, the structure of our anchor bnAb T009-3E04 fit well to the reconstructed nsEM 3D density map of the R996-3C01–HA complex (Figure S6H). This same SWP motif was also found in a human anchor antibody, suggesting that there may be some flexibility within this motif (Figure S6I).

We then compared the structure of T009-3E04 to three known human anchor mAbs, FISW84^19^, 222-1C06^21^ and 204-1B06^20^, and found near identical epitope footprints as well as structural alignment of the CDR H2 and L3 loops across these anchor antibodies (Figure 6E). Comparing the per residue contribution of each antibody further highlights the similarities, including substantial BSA contributions from I/L55_HC_ and Y58_HC_ of CDR H2 as well as the CDR L3 NWP motif (Figure 6F). Importantly, the above key contacts are germline-encoded, suggesting the basis for public usage in humans and macaques.

### H1ssF vaccination elicits equivalent polyclonal serum antibody responses across humans and macaques

Using serum samples from H1ssF vaccinated macaques and human participants enrolled in a phase 1 clinical trial of the H1ssF vaccine (NCT03814720), we performed electron microscopy polyclonal epitope mapping (EMPEM) to evaluate the composition of the polyclonal serum antibody response to HA. Across humans and macaques, we observed three primary polyclonal antibody populations directed to: 1) the central stem, including ‘CR9114-like’ antibodies; 2) the low stem, including ‘MEDI8852-like’ antibodies; and 3) the anchor epitope (Figure 7A-D and S7A-B). Central stem ‘CR9114-like’ serum antibodies were consistently found across all vaccinated macaques and humans (Figure 7A-D and S7A-B). Other shared polyclonal antibody populations included the low stem ‘MEDI8852-like’ response, present in donor 1 and macaques 6974 and R996, and the anchor epitope which was present across all humans and in 3 out of 4 macaques (Figure 7A-D and S7A-B). Additionally, the cryo-EM structure of BB798E-3C07, our macaque central stem bnAb, superimposed well on the central stem 1 density of the EMPEM 3D reconstruction model for BB798E (Figure 7E). Likewise, the macaque anchor bnAb T009-3E04 corresponded well to the anchor density of the T009 EMPEM 3D reconstruction model (Figure 7F). Interestingly, the anti-HA polyclonal response was more heterogeneous in macaques, with animal T009 and BB798E containing antibodies to a low stem 2 epitope, and to the low stem 2 and central stem 3 epitopes, respectively, which were not observed in the three human donor sera (Figures 7B-E). Indeed, some of the macaque polyclonal specificities (including central stem 2 and 3, and low stem 1 and 2) were recapitulated by Fabs isolated from the ‘other HA-stem’ specificity found in the lower HA cluster (Figure 2F, S7C-D). While diverse, these ‘other HA-stem’ mAbs had minimal or no neutralizing activity. Together, polyclonal serum antibody specificities are largely shared between humans and macaques in response to vaccination with HA stem.

**Figure 7.**
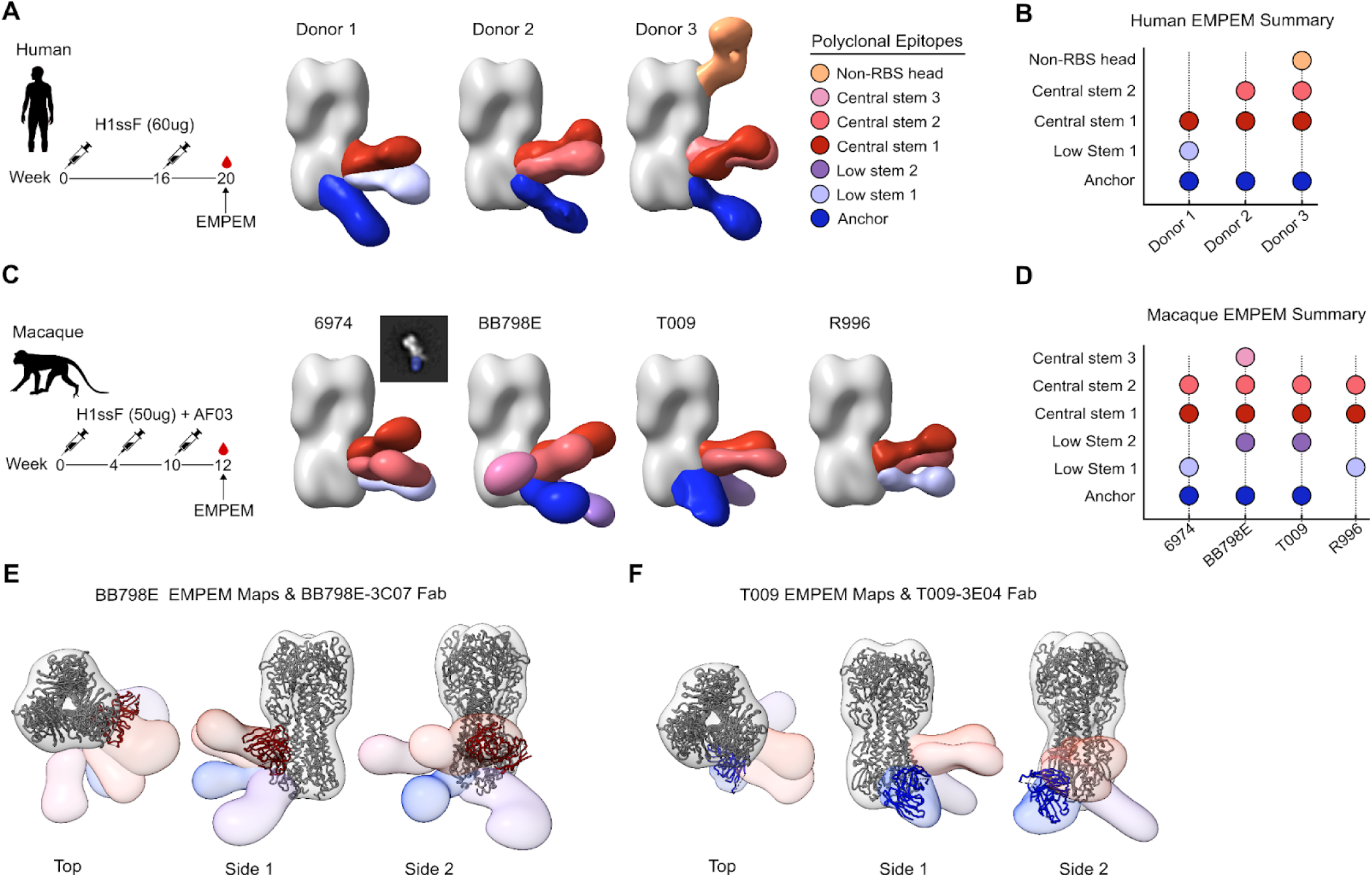
Comparison of polyclonal serum antibody response between humans and macaques vaccinated with H1ssF. **(A-D)** Electron microscopy polyclonal epitope mapping of serum antibody specificities elicited by H1ssF immunization in humans at week 20 (A-B) and macaques at week 12 (C-D). Central stem epitopes include ‘CR9114-like’ serum mAbs and low stem epitopes include ‘MEDI8852-like’ serum mAbs. **(E-F)** Overlays of BB798E-3C07 (central stem bnAb) and T009-3E04 (anchor epitope bnAb) cryoEM bnAb models on polyclonal epitope maps of macaque BB798E (E) and T009 (F), respectively.

## DISCUSSION

Here we demonstrate that humans and macaques can elicit highly convergent bnAb lineages to sites of vulnerability on influenza HA that display similarities at the molecular, immunogenetic, and functional levels. This supports the use of macaques as surrogates for human influenza immunity and underscores the shared evolutionary pathways resulting in convergent public bnAb responses across both species.

Immunoglobulin germline-encoded antigen recognition has been observed against diverse pathogens including influenza virus^39^, HIV-1^40^, hepatitis C virus (HCV)^41^, SARS-CoV-2^42^, and malaria^43^. Such responses are often underscored by recurrent or public use of structural motifs encoded by select immunoglobulin genes with cognate affinity for the target epitope, enabling the elicitation of identical responses across genetically unrelated individuals. However, small animal models such as mice and ferrets are genetically distant, lacking relevant human immunoglobulin genes, and as such, vaccine amplification of public human antibody lineages has largely been restricted to transgenic immunoglobulin gene knock-in mice or mice adoptively transferred with select human bnAb precursors B cells^44,45^. Despite the importance of such models in elucidating principles of vaccine immunity, it remains challenging to recapitulate the complexity associated with the human immune system.

V_H_1-69-containing BCRs represent a major public lineage in human influenza immunity, providing broad activity across group 1 IAVs. While macaques do encode for homologs to V_H_1-69, it was previously thought that such responses were not amenable to elicitation in macaques due to the lack of critical amino acid residues within the antibody paratope. To note, V_H_1-138 usage has been observed in response to influenza infection in macaques^46^, however, detailed characterization of epitope specificity and publicness of such lineages have not been assessed. V_H_1-69 homologs have also been identified in HCV AR3-specific bnAbs in rhesus macaques^47,48^, but has yet to be reported as a predominant public lineage in any non-human primate model for influenza. To date, the only human-like influenza stem bnAb lineage identified most closely resembles that of MEDI8852 (V_H_6-1+D_H_3-3 in humans) which presents as V_H_3/4+D_H_3-41 in macaques^25,26^, a lineage we also find represented within central stem BCRs in this study.

The overwhelming use of V_H_1-138 in central stem-targeting BCRs, with up to 70% of the upper HA^+^ cluster, suggests that the immunoglobulin repertoire in macaques may be similarly endowed with intrinsic affinity to the HA stem. While CDR H2 residues I53/F54 are indispensable in human V_H_1-69 bnAbs^14^, we found that within a macaque V_H_1-138 bnAb, CDR H2 V54 and I56 replace these critical contacts, providing the same germline-encoded ‘masterkey’ function^15,16^. Human V_H_1-69 bnAbs are also marked by low levels of SHM^14,39^ and likewise, we find that macaque V_H_1-138 central stem bnAbs had on average 3.7% SHM, falling below the 4-6% SHM range typically observed in high-affinity human antibodies to pathogens^49^. This occurred using a single immunogen in the absence of prior influenza immunity, contrasting the complex affinity maturation pathways often exhibited by bnAbs to other hypervariable viruses, such as HIV-1^49^.

Canonical anchor antibodies are highly stereotyped with germline encoded contacts, including the CDR H2 Y58_HC_ and CDR L3 NWP motif, which likely contribute to the high degree of structural convergence between macaque and human anchor antibodies. While the NWP-containing lineage is predominant within human anchor antibodies, in macaques, this lineage, although public, was less abundant. Human immunity to influenza is underscored by extensive immune histories which shape the memory repertoire^50^, and it is thought that NWP-containing anchor BCRs may be established by repeated exposures as they are largely derived from class switched and somatically mutated memory B cells and plasmablasts^20,21^. Thus, we cannot exclude the possibility that infrequent anchor antibodies may, in part, be due to the lack of preexisting immunity in our animals.

Advances in computational and structural immunogen design have enabled the development of germline-targeting immunogens to activate and expand functional public human bnAb lineages with atomic level accuracy^44,51^. However, the utility of macaques to tackle human immunological challenges has remained a longstanding question. Recent attempts to elicit anti-HIV-1 Env CD4-binding site and BG18-like bnAb precursors in macaques have highlighted convergent bnAb development pathways driven by human-like encoded motifs^52,53^. These studies as well as ours have emphasized the immunological relevance of macaques to humans while also providing a framework to experimentally address questions that may not be feasible in human clinical trials.

At a broader level, convergence in public B cell immunity across macaques and humans is intriguing given the diversity in their immunoglobulin repertoires. Despite diverging from a common ancestor ∼25 million years ago^24^, the utility of homologous public immunoglobulin germline-encoded motifs with specificity to two distinct HA stem supersites suggests that the repertoires of both species may have been shaped by similar evolutionary pressures. How, then, might identical germline-encoded components of the BCR be selected for and maintained in both repertoires? While the antibody repertoires of humans and macaques were shaped by positive selection for survival advantage, we cannot ascribe selection of a specific immunoglobulin gene to a single pathogen. For example, biased usage of V_H_1-69 is also found in response to HIV-1^54^, HCV^41^, SARS-CoV-2^55^, and malaria^56^. Rather the primary role of the immune system is to ensure survival from infection and it has long been proposed that such germline-encoded structural motifs may function as an ‘innate-like’ pattern recognition element, responsible for mounting rapid and robust defenses against antigenic structures on pathogens^57–59^. Indeed, we find that V_H_1-69 homologs are preserved across the primate order, suggesting widespread utility of this V_H_-gene. While we can only speculate that germline-encoded affinity may have originally been selected as an ‘SOS’ component in the antibody repertoires of humans and macaques^57^, these responses have likely been co-opted for secondary use as influenza HA stem bnAb responses, resulting in their stereotyped usage across both species^58^.

## LIMITATIONS OF THE STUDY

The advancement of sequencing technologies has greatly improved assignment of macaque immunoglobulin genes; however, classification of such genes and their corresponding alleles remain incomplete. This will undoubtedly influence the accuracy of SHM measurement and allelic assignment of our macaque BCR genes. Additionally, as mentioned above, macaques in this study lack pre-existing influenza immunity, a parameter that, while important, will be difficult to reconstitute experimentally in macaques. As such, we do not assess how such pre-exposure alters the anti-influenza antibody landscape, particularly in regard to eliciting central stem and anchor B cells in macaques. However, the fact that we identify equivalent central stem and anchor lineages in this study suggests that despite lacking this physiologically relevant variable, macaques are still amenable to elicitation of such responses.

## METHODS

### Cynomolgus Macaque PBMCs

PBMC samples from cynomolgus macaques (*Macaca fascicularis*) were collected from a historic HA stem nanoparticle (H1ssF) vaccination study^25^. All experimental procedures, protocols and care of the animals were approved by the Institutional Animal Care and Use Committee at the Vaccine Research Center, NIH or Bioqual. Animals were used in studies in accordance with all federal regulations, NIH guidelines, AAALAC, and IACUC approval.

### Mice

Female Balb/cJ mice, 6-8 weeks old, were purchased commercially from Jackson Laboratories (Bar Harbor, ME). Upon arrival, mice were housed in standard microisolator caging at ABSL-2, five per cage. All animal procedures were performed at the University of Pittsburgh under the approval of the Institutional Animal Care and Use Committee, protocol 22040682. Animal care and use was in accordance with the Guide for the Care and Use of Laboratory Animals of the National Research Council and with the Association for Assessment and Accreditation of Laboratory Animal Care (AAALAC).

### Cell Lines

Expi293 cells (ThermoFisher Scientific) were cultured in Expi293 Expression Medium (Life Technologies at 37°C with 8% CO_2_ and agitation at 120 rpm. MDCK-SIAT1-PB1 cells used for microneutralization have been described previously^60^.

### Human Serum Samples

Human serum samples were taken from the H1ssF phase 1 clinical trial (ClinicalTrials.gov identifier NCT03814720), where healthy adults (aged 18 to 70) were given 60 µg H1ssF twice 16 weeks apart^23^. Samples used in this study were collected at week 20 (see Figure 7A). Trial protocols were reviewed and approved by the NIAID Institutional Review Board. Informed consent was obtained from every enrolled participant and complied with all relevant ethical regulations. Compensation was provided for time and effort related to participation in the clinical trial.

### Expression and Purification of Recombinant HAs

Recombinant soluble full length and stem-only HA trimers are all derivations H1N1 of A/New Caledonia/20/1999 (H1 NC99) and have been described previously^13,21,28^. The H1-Δcentral and H1-Δanchor HA probes were modified to display glycans at I45N_HA2_ (H1-Δcentral) or at positions Q27N_HA2_ and N29T_HA2_ (H1-Δanchor). All full-length soluble HA trimers contain the Y98F mutation within the RBD to prevent sialic acid binding^61^. Expression vectors encoding HA constructs were transfected into Expi293 cells at 1 ug/ml using the Expifectamine293 transfection kit (ThermoFisher) according to manufacturer’s instructions. Following a 4-day incubation period, proteins were harvested from culture supernatant, filtered, and purified using Ni-Sepharose excel beads (Cytiva) by gravity flow. The beads were washed with 50 mM Tris-HCl pH 8, 0.5M NaCl containing 30 mM imidazole and then eluted with buffer containing 300 mM imidazole. HA trimers were purified via size exclusion chromatography (SEC) using the AKTA pure protein purification system and Superdex 200 10/300 (Cytiva) column. Antigenicity of all recombinant HA proteins were confirmed by binding to the conformational antibodies 315-02-1H01, FISW84, and CH65. DS-Cav1 (RSV-F) protein was purified as described above using Ni-Sepharose excel beads.

Stem-only HA trimers were biotinylated via the AviTag using biotin protein ligase (Avidity) based on manufacturer’s instructions. Biotinylated HAs were conjugated with fluorescently labeled streptavidin at a 4:1 molar ratio to generate fluorescently labeled HA tetramers. Full length HA trimers were directly labeled according to manufacturer’s instructions (ThermoFisher Scientific, A30006).

### Production of Monoclonal Antibodies and Fabs

Heavy and light chain variable regions from monoclonal antibodies generated in this study were synthesized as gene blocks (GenScript) and cloned into expression vectors to human IgG1 heavy and either kappa or lambda light chains. Monoclonal antibodies were recombinantly produced by transfection of heavy and light chain DNA into Expi293 cells using ExpiFectamine. Following 4 days of incubation, the antibodies were purified from cell culture supernatant using Protein A resin (GE Healthcare) and eluted with IgG elution buffer into 100mM Tris, 300mM NaCl, pH 8.0 and buffer exchanged into PBS.

For Fab production and purification, monoclonal IgG were cleaved with endoproteinase Lys-C (Promega) overnight at room temperature. Upon completion, the digestion was quenched using protease inhibitor (Roche) and Fab components were separated by passing through protein A or protein L columns.

### Flow Cytometry and B Cell Sorting

Cryopreserved peripheral blood mononuclear cells (PBMCs) were stained on ice with a cocktail of the following antibodies: CD16 BV510 (clone 3G8), CD3 BV510 (clone SP34-2), CD56 BV510 (clone B159), CD14 BV510 (clone M5E2), CD20 APC/Cyanine7 (clone 2H7), IgM PerCP-Cy5.5 (clone G20-127), IgG BUV395 (clone G18-145). To isolate vaccine elicited HA stem specific B cells, the following cocktail also contained H1 NC99 probes including stem-only HA labeled with streptavidin-phycoerythrin (H1-stem PE) and full-length HA directly labeled with AF488 (H1-HA AF488). The full length HA probe contained the Y98F mutation within the RBD which prevents surface sialic acid binding. All recombinant HA proteins were quality controlled by size exclusion chromatography (SEC) and by binding to conformational antibodies 315-02-1H01, CR9114, CH65, and FISW84. Live/dead fixable Blue (ThermoFisher Scientific) was used to assess viability.

Live cells were sorted based on reactivity to CD20^+^/CD3^-^/CD14^-^/CD16^-^/CD56^-^/IgM^-^/IgG^+^/H1-HA^+^/H1-stem^+^ into 96 well plates containing lysis buffer (RLT Buffer with 1% 2-mercaptoethanol) and immediately frozen at –80°C. Index sorting was applied to assign specificity of ‘upper’ and ‘lower’ clusters. All samples were analyzed on the FACS Aria II (BD Biosciences) and downstream data analysis was performed using FlowJo software version 10.8.2 (TreeStar).

### Rapid assembly, transfection, and production of immunoglobulins (RATP-Ig)

To sequence single BCRs and to screen antigen specific B cells, Rapid assembly, transfection, and production of immunoglobulins (RATP-Ig) was applied as described previously^29^. Briefly, RNA from single cell sorted B cells were purified and cDNA was synthesized by 5’ RACE using a modified version of the SMARTSeq-V4 protocol. Heavy and light chain variable regions were enriched from cDNA using IgG or IgK/IgL primer pools and sequenced using paired end 2×150 reads on an Illumina MiSeq system. Heavy and light chain sequence pairs were visualized using Circos^62^. Sequence motifs were generated using WebLogo.

For production of immunoglobulins, the enriched heavy and light chain variable region fragments were assembled into a single linear cassette. Each cassette contained a CMV promoter and a TBGH poly A fragment. Linear cassettes encoding the monoclonal heavy and light chain genes were further amplified by PCR and transfected into Expi293 cells in 96-well deep-well plates using the Expi293 Expression System (ThermoFisher Scientific). Cell cultures were incubated at 37°C and 8% CO_2_, with shaking at 1100 RPM for 5–7 days. Culture supernatants were clarified by centrifugation.

## ELISA

HA protein antigens (H1-HA, H1-Δcentral, H1-Δanchor, H1-Δcentral/Δanchor) were coated onto 96 well Nunc MaxiSorp plates at 200 ng HA per well and incubated overnight at 4°C. Plates were blocked with 5% milk in PBS for 1 h and washed with PBS and 0.05% Tween 20 (PBST). The plates were incubated for 1 h with either immune serum (starting at a 1:100 dilution) or monoclonal antibodies (starting at 10 µg/ml) and were diluted down 5-fold in PBS + 5% milk. RATP-Ig supernatants were tested at a 1:5 dilution in PBS + 5% milk. ELISA plates were washed and incubated with mouse anti-monkey IgG-HRP (Southern Biotech) or goat anti-human IgG-HRP (Southern Biotech) antibodies at 1:5000 dilution for 1 h. Plates were again washed and developed with TMB substrate and quenched with 1 N sulphuric acid (H_2_SO_4_) and read at 450 nm using the Spectramax plus 384 microplate reader (Molecular Devices).

### Microneutralization Assays

The microneutralization assay and reporter viruses used have been previously described^60^. Briefly, all viruses used were generated with either modified PB1segments expressing the TdKatushka reporter gene (R3ΔPB1), including A/New Caledonia/20/1999 (H1N1), A/Michigan/45/2015 (H1N1) and A/Wisconsin/67/2005 (H3N2). For H5N1 and H2N2 viruses, both PB1 and HA segments were modified (‘rewired’) to prevent HA reassortment: the PB1 segment encodes the HA coding region whereas the HA segment encodes the TdKatushka reporter gene (R4ΔPB1). Monoclonal antibodies were tested at a starting concentration of either 25 µg/ml and serially diluted four-fold. Antibodies were incubated for 1 h at 37°C with pre-titrated A/New Caledonia/20/1999 (H1N1), A/Michigan/45/2015 (H1N1), A/Vietnam/1203/2004 (H5N1), A/Singapore/1/1957 (H2N2), or A/Wisconsin/67/2005 (H3N2) viruses. Virus and antibody mixtures were transferred to 96 well plates (PerkinElmer) and mixed with 1.0 × 10^4^ MDCK-SIAT1-PB1 cells and incubated overnight at 37°C. The following day, the number of fluorescent cells in each well was counted using the Celigo image cytometer (Nexcelom Biosciences). IC_50_ titers were calculated in Prism.

For serum microneutralization, serum samples were treated with receptor destroying enzyme (RDE II, Denka Seiken) at 1 part serum to 3 parts RDE, incubated overnight at 37°C and heat inactivated at 56°C. Serum samples were tested at an initial dilution of 1:40 and diluted down. For the competition reporter microneutralization assay, RDE treated NHP sera was serially diluted as described above and incubated with 0.1 mg of either H1-HA, H1-Δcentral, H1-Δanchor and DS-Cav1 for 1 h at RT before addition of virus and cells.

### Assessment of Monoclonal Antibody Autoreactivity

Antibody autoreactivity was assessed by ANA HEp-2 staining (ZEUS Scientific, reference number FA2400EB) and anti-cardiolipin ELISA (Inova Diagnostics, reference number 708625). Anti-HIV-1 antibodies VRC01LS, 4E10, VRC07-523LS and VRC07-523 G54W were used as controls^37^. For the ANA HEp-2 staining, all antibodies were tested at 25 and 50 μg/mL following manufacturer’s instructions. Images were taken on a Nikon Ts2R microscope for 500 ms using a 20 × lens. VRC01LS, 4E10, VRC07-523LS and VRC07-G54W were defined with scores from 0 to 3. Test antibodies were scored by visual estimation of fluorescent intensity resulting from their binding to HEp-2 cells compared to those of the control antibodies. Scores equal to or greater than 1 at 25 μg/mL were classified as autoreactive and between 0 and 1 as mildly autoreactive. For the anti-cardiolipin ELISA, antibodies were tested at a starting concentration of 100 μg/mL followed by 3-fold serial dilutions. IgG phospholipid (GPL) units were derived from the standard curve. GPL score < 20 was considered as not reactive, 20–80 as low positive and > 80 as high positive.

### Passive Transfer and Viral Challenge

Antibodies were given by intraperitoneal inoculation, 200 µl given by a 1 cc syringe and 25-gauge needle. Virus challenge was by intranasal inoculation using 50 µl applied by micropipette to the nares while mice were anesthetized. Following viral challenge, mice were weighed daily and checked twice daily for changes in clinical signs indicative of morbidity (changes in appearance and behavior). Mice that were either moribund, suffering respiratory distress, a > 20% loss in body weight from baseline, or hindlimb paralysis were euthanized immediately. Euthanasia was performed using procedures consistent with American Veterinary Medical Association guidelines.

Mouse-adapted A/California/07/2009 was propagated in chicken eggs to produce a large stock which was aliquoted and frozen for all studies. The virus concentration (3.75 × 10^6^ pfu/ml) was titered by plaque assay prior to use to determine stock concentration. Preliminary studies found no differences in survival (or survival time) between female and male mice and established an LD_50_ of 5,300 pfu. Mice were challenged with 8 × 10^4^ pfu (15 LD_50_). Dose was confirmed by plaque assay. An infectious clone of A/Vietnam/1203/2004 (H5N1) was propagated in chicken eggs to produce a large stock which was aliquoted and frozen for all studies. The virus concentration (2 × 10^9^ pfu/ml) was titered by plaque assay prior to use to determine stock concentration. Preliminary studies established an LD_50_ of 2 pfu. Mice were challenged with 12 pfu (6 × LD_50_). Dose was confirmed by plaque assay curve.

### Nonhuman Primate IGHV1-69 Phylogenetic Tree

Using the Ensemble software, we took the known human alleles for *IGHV1-69* and performed a BLAST analysis against the available genome sequences. Taxonomic identifiers were extracted from the NCBI taxonomy browser. The phylogenetic tree was generated using NCBI tree elements in PhyloT and then visualized using Interactive Tree of Life (iTOL). The degree of nonhuman primate homology to human IGHV1-69*01 was evaluated at the nucleotide level using Clustal Omega.

### BLI

BLI experiments were performed using the Octet HTX instrument (ForeBio). All biosensors were hydrated in PBS prior to use. Ni-NTA biosensors (ForteBio) were loaded with his-tagged recombinant H1 NC99 HA. After equilibration for 60 s in assay buffer (25 mM Tris, 150 mM NaCL, 1% BSA, pH 8.0), biosensors were dipped in either Fabs (30 µg/ml) for mAbs (30 µg/ml) for 600 s, followed by dissociation for 600 s. All assay steps were performed at 30°C with shaking at 1000 RPM.

### Cryo-EM sample preparation, data collection and structure determination

To assemble the HA-Fab complex, HA NC99 and Fab BB798E-3C07 or Fab T009-3E04 were mixed at a molar ratio of 1:1.5 (HA protomer:Fab) and incubated for 5 min at room temperature. Graphene covered grids were produced in-house by following a published protocol^63^ based on Quantifoil R 2/2 gold grids. Graphene grids were treated with ozone using a HELIOS-500 UVFAB UV-ozone cleaner for 10 min immediately before use. Vitrification was performed at a protein concentration of 0.01 mg/ml using a Thermo Scientific Vitrobot Mark IV plunger with the following parameters: sample volume of 2.7 µl, chamber humidity of 95% and chamber temperature of 4°C. A total of 10,544 movies for the HA NC99/Fab BB798E-3C07 complex and 11,390 movies for the HA NC99/Fab T009-3E04 complex were collected on an FEI Titan Krios G1 electron microscope equipped with a Gatan K2 Summit direct electron detector operated in the counting mode (Table S1).

For the HA NC99/Fab BB798E-3C07 complex dataset, single particle analysis was performed in cryoSPARC 4.3^64^ (Figures S5A-S5G). After patch motion correction and patch contrast transfer function (CTF) parameters estimation, 9,591 micrographs were selected for downstream processing based on full-frame motion, CTF fit resolution and relative ice thickness. Particles were picked from a subset of 100 micrographs using the blob picker and, after several rounds of 2D classification, used for Topaz training^65^. The resulting Topaz model was used for particle picking in the entire dataset. After multiple rounds of 2D classification, two independent runs of *ab initio* reconstruction were performed, followed by heterogeneous refinement of the particle subsets producing complete, high-resolution maps of the complex. The two subsets were then merged, followed by removal of duplicates. The resulting 104,541 particles were subjected to non-uniform refinement^66^ to generate the final map.

For the HA NC99/Fab T009-3E04 complex dataset, cryoSPARC 4.3 and Relion 4.0^67^ were used to obtain the 3D map (Figure S5H - S5N). Movie frames were aligned with MotionCor2^68^, and CTF parameters were estimated using ctffind4^69^. 10,025 micrographs with a CTF fit resolution of at least 5 Å were used for further processing. Templates for template-based automatic particle picking in Relion were obtained by 2D classification of particles picked from 200 micrographs using the Laplacian-of-Gaussian algorithm. This subset was also used for Topaz training. The particles from template picking and Topaz picking were merged, followed by elimination of duplicates. The subsequent processing steps were similar to those performed for the HA NC99/Fab BB798E-3C07 complex, except that the final particle stack produced by heterogeneous refinement in cryoSPARC was subjected to particle polishing in RELION. The resulting 160,012 particles were used for non-uniform refinement in cryoSPARC to generate the final map. The reported resolutions were determined based on the “gold standard” criterion at the FSC curve threshold 0.143^70^. Local resolution was estimated using ResMap^71^.

To obtain the atomic models of the complexes, a previously deposited hemagglutinin NC99 structure from PDB entry 8D21 was docked into the cryo-EM density, along with the initial models of the Fab generated with ColabFold^72^, using UCSF Chimera^73^. The atomic models were refined by alternating rounds of real-space refinement in Phenix^74^ and model building in Coot^75^ and ISOLDE^76^. Molprobity was used to validate the final models^77^. Map-model correlation was assessed with phenix.mtriage using the FSC curve threshold of 0.5^78^. The refinement statistics are summarized in Table S1.

### Negative-stain Electron Microscopy

To prepare complexes, HA was mixed with Fab at a slight molar excess of the latter in buffer composed of 10 mM HEPES, pH 7.0, and 150 mM NaCl to a final protein concentration of about 0.02 mg/ml. The sample was applied to a freshly glow discharged carbon-coated copper grid for about 15 s, and excess liquid was removed with blotting paper. The grid was washed two times with the above buffer, followed by negative staining with 0.75% uranyl formate. Data was collected using a TermoFisher Talos F200C electron microscope operated at 200 kV and equipped with a Ceta camera at a nominal magnification of 57,000 (corresponding to a pixel size of 2.53 Å). Topaz was used for particle picking^65^. The following steps included 2D classification, 3D classification and 2D refinement, which were performed in Relion467. 3D maps were visualized and aligned using UCSF Chimera^73^.

### EMPEM

EMPEM protocols were performed as described previously^79^. Sera was heat inactivated in a 56°C water bath for 60 min and incubated on CaptureSelect IgG-Fc resin (Thermo Scientific) in a 1:1 volume ratio of serum to resin overnight. IgG-depleted sera was removed and the resin was washed three times with 5 column volumes (CV) of 1 × PBS. IgG was eluted by incubation with 10 CV 0.1M glycine pH 2.0 buffer for 20 min followed by neutralization with 1M Tris-HCl pH 8 buffer, repeated twice. Samples were buffer exchanged into 1 × PBS using centrifugation with Amicon concentrators. Total IgG was digested with activated papain in digestion buffer (20 mM sodium phosphate, 10 mM EDTA, 20 mM cysteine, pH 7.4) at 37°C for 5 h and was quenched with Iodacetamide. Samples were buffer exchanged into 1 × PBS. To purify any further serum impurities and papain, the polyclonal fab mixture was run on size exclusion chromatography with a Superdex 200 increase 10/300 column (“S200i,” GE Healthcare) in 1 × TBS buffer. Fab rich fractions were pooled and concentrated for EMPEM complexing. 0.5 mg of each polyclonal Fab mixture was mixed with 10 µg HA, incubated at room temperature overnight, and purified over an S200i column. The SEC peak corresponding to the polyclonal complex was collected, concentrated, immediately added to a negative stain electron microscope (nsEM) grid, and stained with 2% uranyl formate. Polyclonal complexes were imaged on a Tecnai Spirit electron microscope at a nominal magnification of 52,000 ×, a pixel size of 2.06 Å, a defocus value of –1.5 µm, and an electron dose was 25 e^-^/Å^2^. Micrographs were recorded using a Tietz (4k) TemCam-F416 CMOS. Automated acquisition was performed with Leginon^80^ and processed on Appion^81^. DoG Picker^82^ was used to choose particles and 2D and 3D classification was performed in Relion version 3.0^83^. Figures were made with UCSF ChimeraX^73^.

## QUANTIFICATION AND STATISTICAL ANALYSIS

All statistical analyses were conducted using Graphpad Prism software. Sample sizes and statistical tests are indicated in the figure legends.

## RESOURCE AVAILABILITY

### Lead contact

Requests for further information and resources should be directed to and will be fulfilled by the lead contact, Masaru Kanekiyo.

### Materials availability

All unique antibodies and other reagents generated in this study will be made available on request from the lead contact with a completed materials transfer agreement.

### Data and code availability

The PDB and EMDB accession numbers for BB798E-3C07 Fab-HA and T009-3E04 Fab-HA complexes reported in this paper are 9CJY/EMD-45636 and 9CJZ/EMD-45637, respectively. Accession IDs for EMPEM are EMD-46824, EMD-46825, EMD-46827, EMD-46829, EMD-46830, EMD-46831, EMD-46832.

## ACKNOWLEDGEMENTS

The authors thank L. Dropulic and the VRC 321 study team and study participants for human samples; **(A)** G. Nabel, N. Darricarrere, and C.J. Wei (formerly Sanofi) for cryopreserved macaque PBMC samples. **(B)** D. Ambrozak (VRC) for assistance with cell sorting; D. Lingwood (Harvard) and members of the VRC Influenza Program for helpful discussion. This work was supported, in part, by the Vaccine Research Center, an intramural Division of National Institute of Allergy and Infectious Diseases, National Institutes of Health; federal funds from the Frederick National Laboratory for Cancer Research, National Institutes of Health, under contract HHSN261200800001 (Y.T and N.L.). J.H. and A.B.W. are supported by CIVIC 75N93019C00051. M.C. and G.B.K.H. are funded by a grant from the Swedish Research Council (2017-00968). The content is solely the responsibility of the authors and does not necessarily represent the official view of the National Institutes of Health.

## AUTHOR CONTRIBUTIONS

Conceptualization: M.S and M.K.; formal analysis: M.S., Y.T., J.H., G.K.H., C.S.; investigation: M.S., N.L., Y.T., M.D.R., J.H., A.J.R.., J.A.F., A.R.H., S.C.S., J. RT., C.L., J.S.M., T.S., C.W., E.M., M.C., S.F.A.; resources: A.C, M.R.; writing - original draft preparation: M.S., M.K.; writing - review and editing: all authors; supervision: T.C.P., G.B.H.K., C.A.S., D.S.R., D.C.D., T.Z., A.B.W., M.K.; project administration: R.A.G.; funding acquisition, T.C.P., G.B.K.H., D.S.R., D.C.D., T.Z., A.B.W., M.K.

## DECLARATION OF INTERESTS

M.K. is named as an inventor on patents describing the HA stem immunogen used in this study under US10363301, US11147867, US11679151 which were filed by the Department of Health and Human Services.

## SUPPLEMENTAL INFORMATION

### Supplemental information

**Figure S1.**
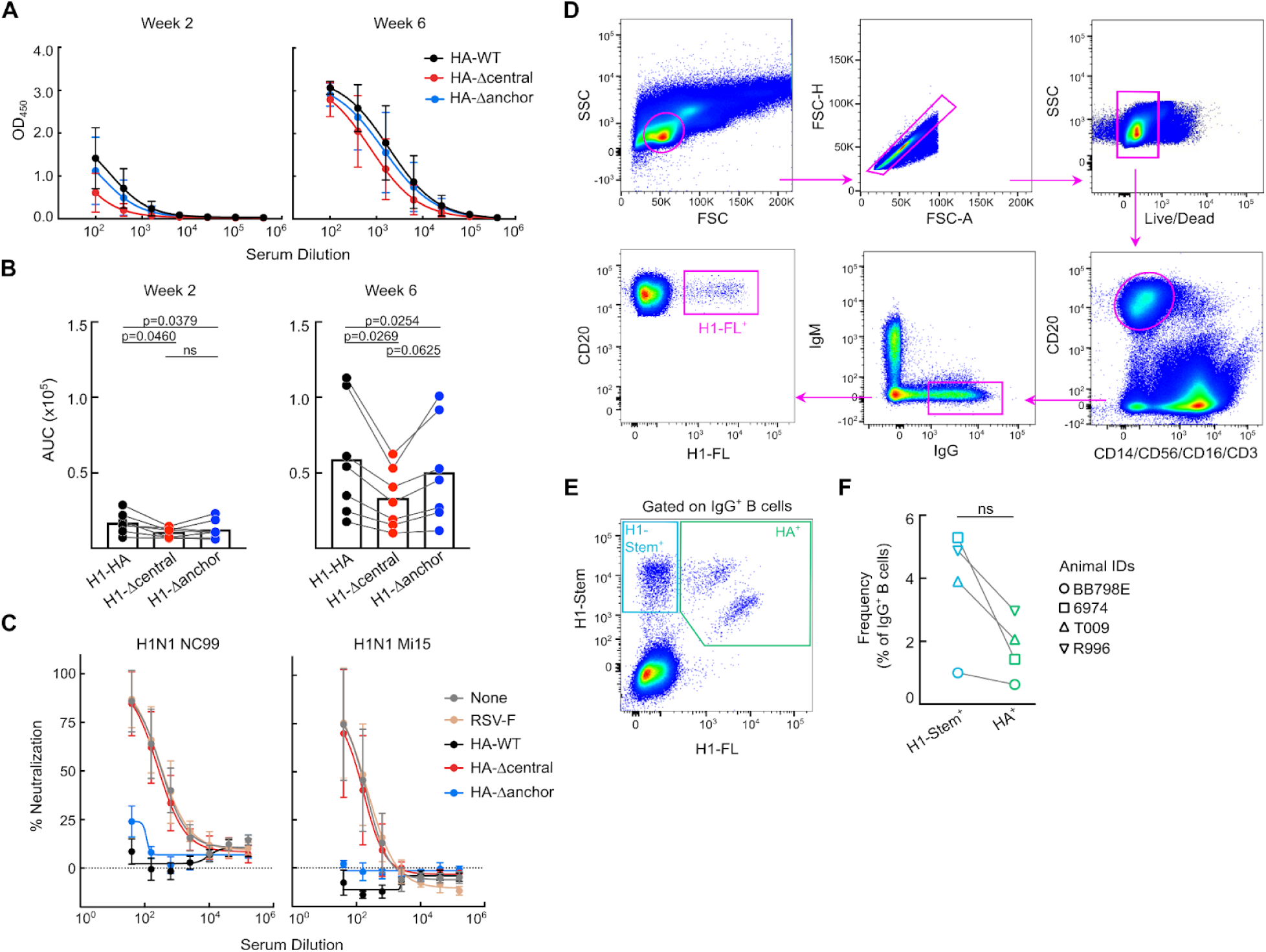
Detailed characteristics of macaque immune responses, related to Figures 1 and 2. **(A-B)** Serum reactivity to HA-WT, HA-Δcentral and HA-Δanchor soluble probes at weeks 2 and week **(A)** 6. Shown are mean and SD (A). Corresponding AUC at weeks 2 and 6. Statistical significance determined by one-way ANOVA with Geisser-Greenhouse correction and Tukeys post hoc test (B). **(B)** Serum neutralization curves following serum pre-absorption with RSV-F, HA-WT, HA-Δcentral and HA-Δanchor proteins. Neutralization was measured against H1N1 NC99 and H1N1 MI15 reporter viruses. **(C)** Gating strategy to identify antigen specific HA-FL^+^ B cells. PBMCs were gated on CD3^-^/CD14^-^/CD56^-^/CD16^-^/CD20^+^/IgG^+^/IgM^-^/H1-FL^+^. **(E-F)** Representative gating strategy to delineate H1-stem single positive and HA+ B cells (gated on IgG^+^ B cells) (E). Quantification of H1-stem^+^ (light blue) and HA^+^ (green) B cells. Significance determined by two tailed Wilcoxon matched-pairs signed rank test (F).

**Figure S2.**
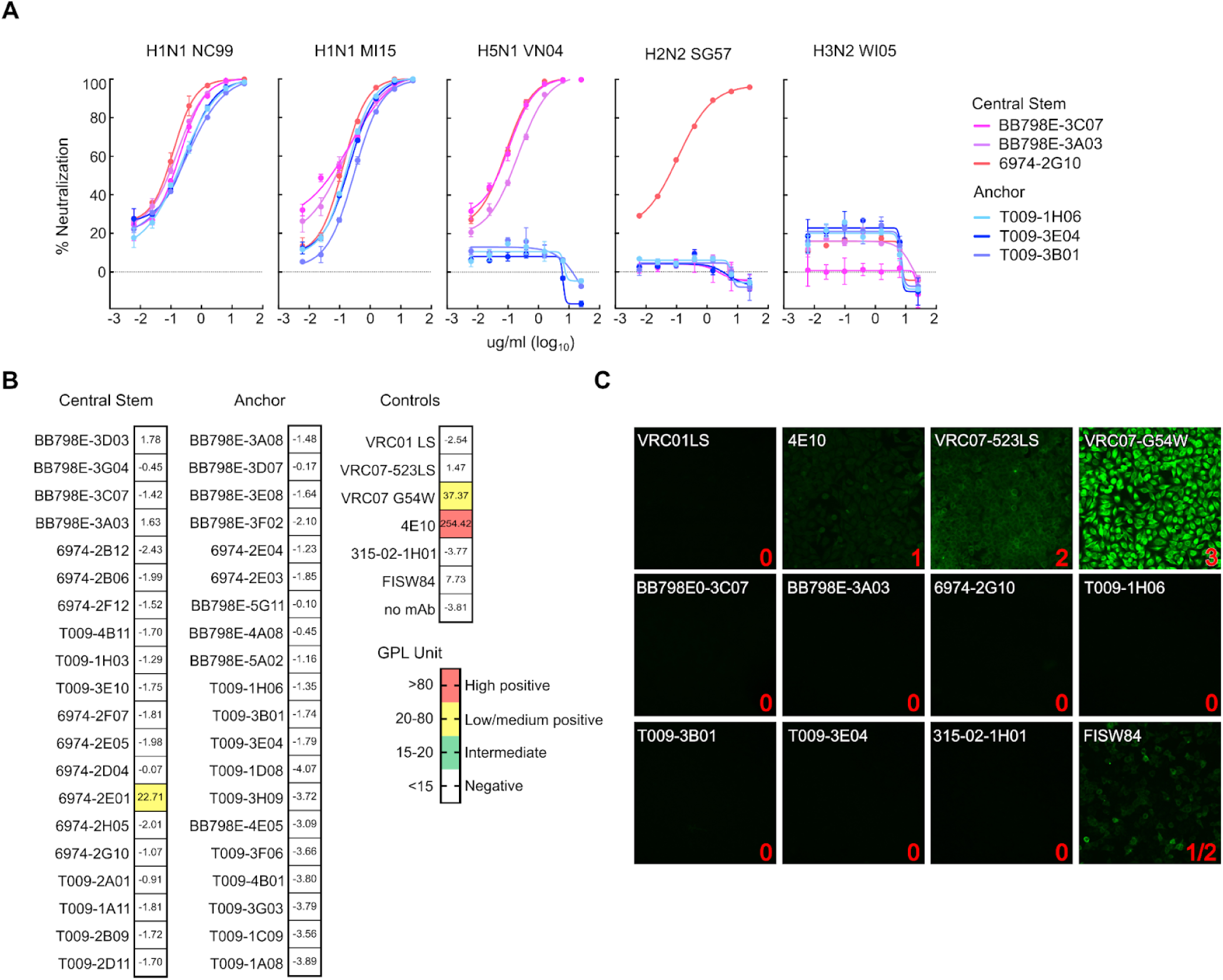
Neutralization and polyreactivity of H1ssF elicited monoclonal antibodies, related to Figure 3. **(A)** Neutralization dilution curves for mAbs BB798E-3C07, BB798E-3A03, 6974-2G10, T009-1H06, T009-3E04, T009-3B01 against H1N1 NC99, H1N1 MI15, H5N1 VN04, H2N2 SG57 and H3N2 WI05 reporter viruses. Shown are mean and SD. **(B)** IgG phospholipid (GPL) unit value for central stem, anchor epitope and control monoclonal antibodies. GPL score < 20 was considered as not reactive, 20–80 as low positive and > 80 as high positive. **(C)** ANA HEp-2 staining of select candidate mAbs: BB798E-3C07, BB798E-3A03, 6974-2G10, T009-1H06, T009-3B01, T009-3E04, compared to control mAbs: 315-02-1H01, FISW84, VRC01LS, 4E10, VRC07-523LS, and VRC07 G54W. Binding to Hep2 cells is scored and indicated (numbered 0 - 3; score 1/2 indicates reactivity between 4E10 and VRC07-523LS).

**Figure S3.**
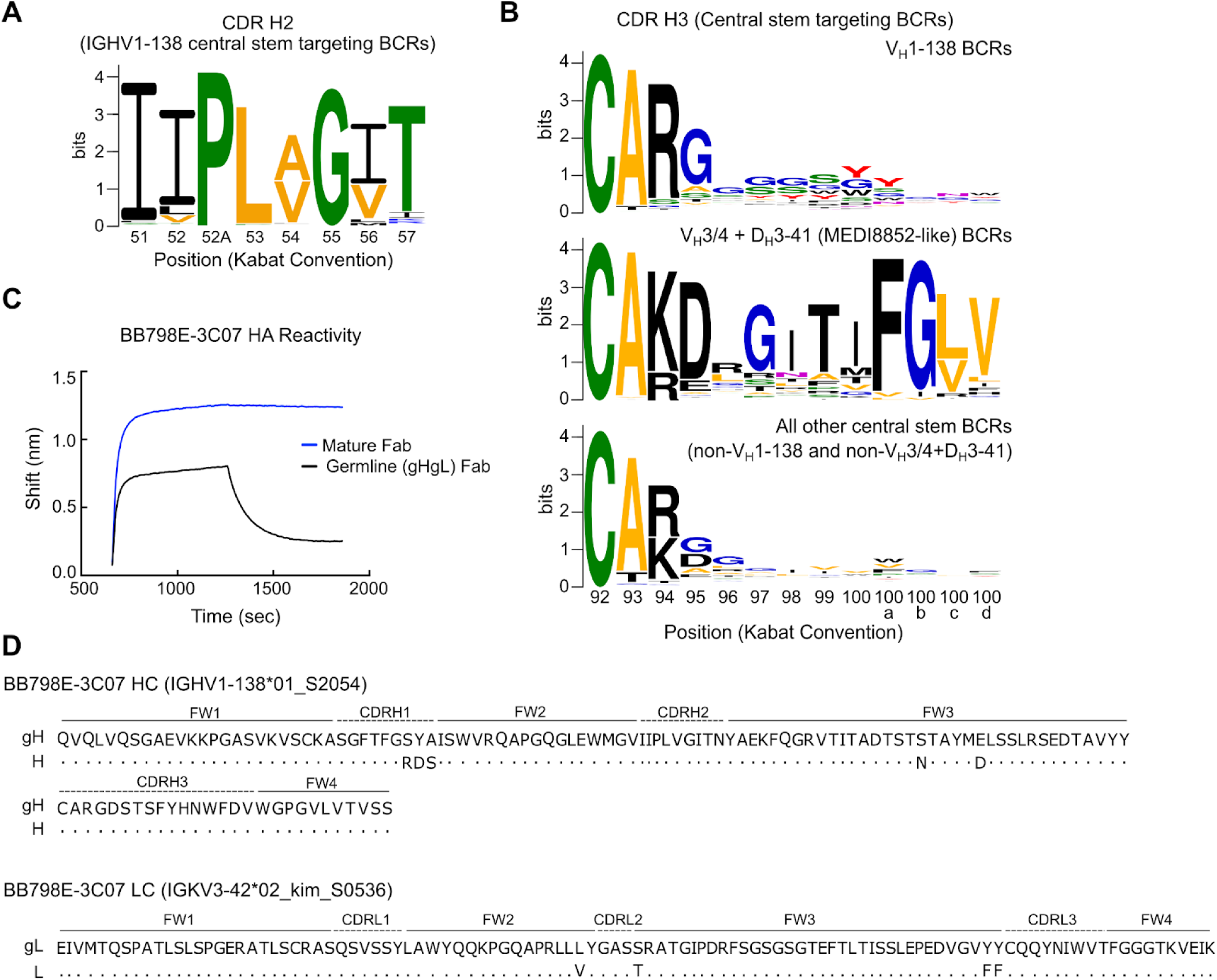
IGHV1-138 central stem mAb characterization, related to Figure 4. **(A)** CDR H2 sequence motif of V_H_1-138 central stem targeting BCRs from 4 macaques. Position indicated by Kabat convention. **(B)** CDR H3 sequence motifs of central stem targeting lineages including V_H_1-138 BCRs (top), V_H_3/4 + D_H_3-41 (MEDI8852-like, middle), and all other central stem targeting BCRs (bottom) from 4 macaques. Tyrosines are colored in red. **(C)** BLI sensogram of gHgL (germline) and mature BB798E-3C07 Fab binding to H1 NC99 HA. **(D)** Amino acid sequence alignment of mature heavy (H) and light (L) chains and germline reverted heavy (gH) and light (gL) chains of BB798E-3C07.

**Figure S4.**
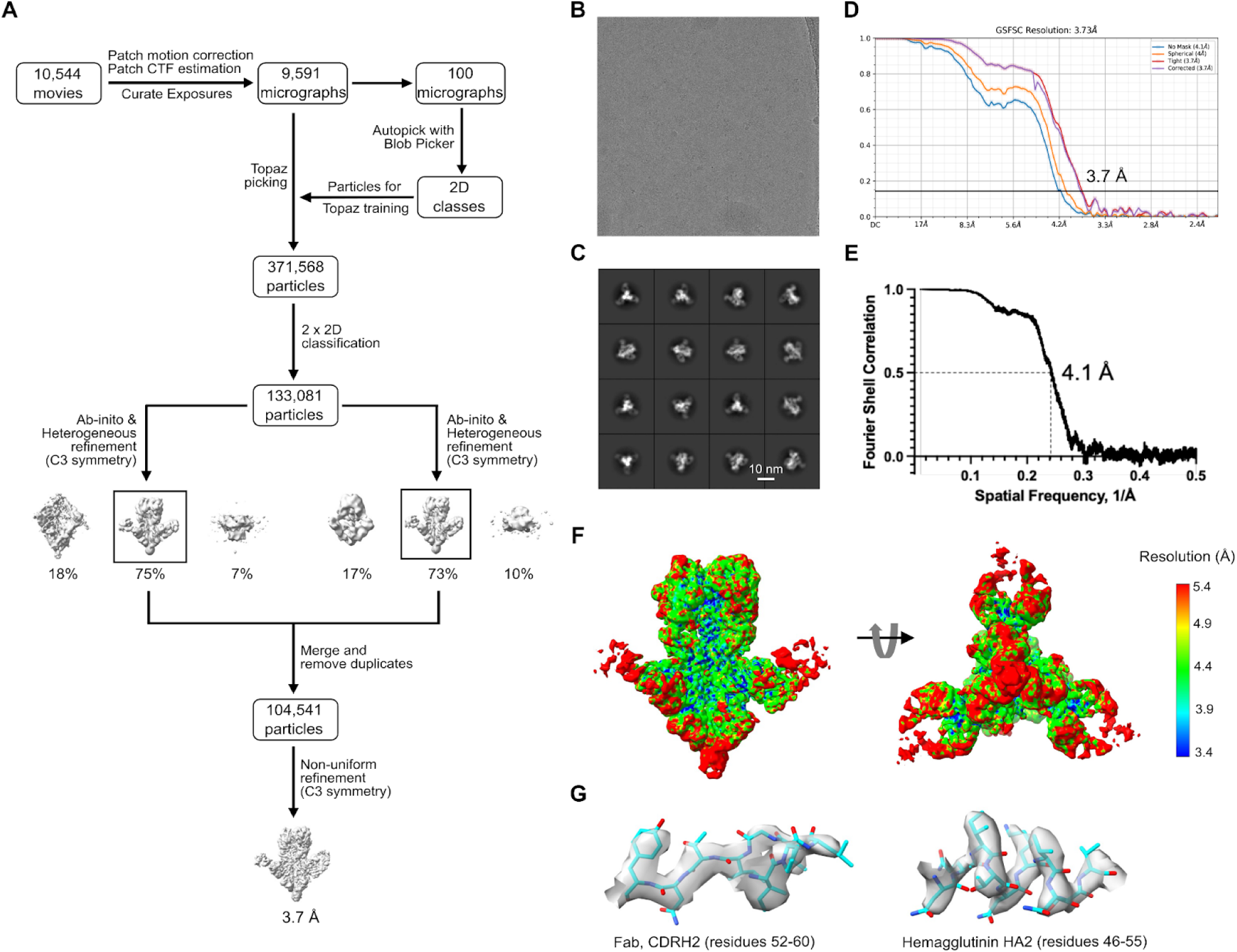
BB798E-3C07 Cryo-EM data processing and workflow, related to Figure 4. **(A)** Cryo-EM data processing workflow for HA NC99 in complex with Fab BB798E 3C07. **(B-C)** Representative micrographs of 3C07 (B), and representative high-resolution 2D class averages for 3C07 (C). **(D)** Gold-standard resolution data generated by cryoSPARC. At the 0.143 threshold, the resolution is 3.7 Å for 3E04. **(E)** Fourier shell correlation curve between the map and the atomic model for 3C07. **(F)** Results of local resolution analysis using ResMap. The cryo-EM map is colored according to local resolution for 3C07. **(G)** Examples of cryo-EM density for 3C07.

**Figure S5.**
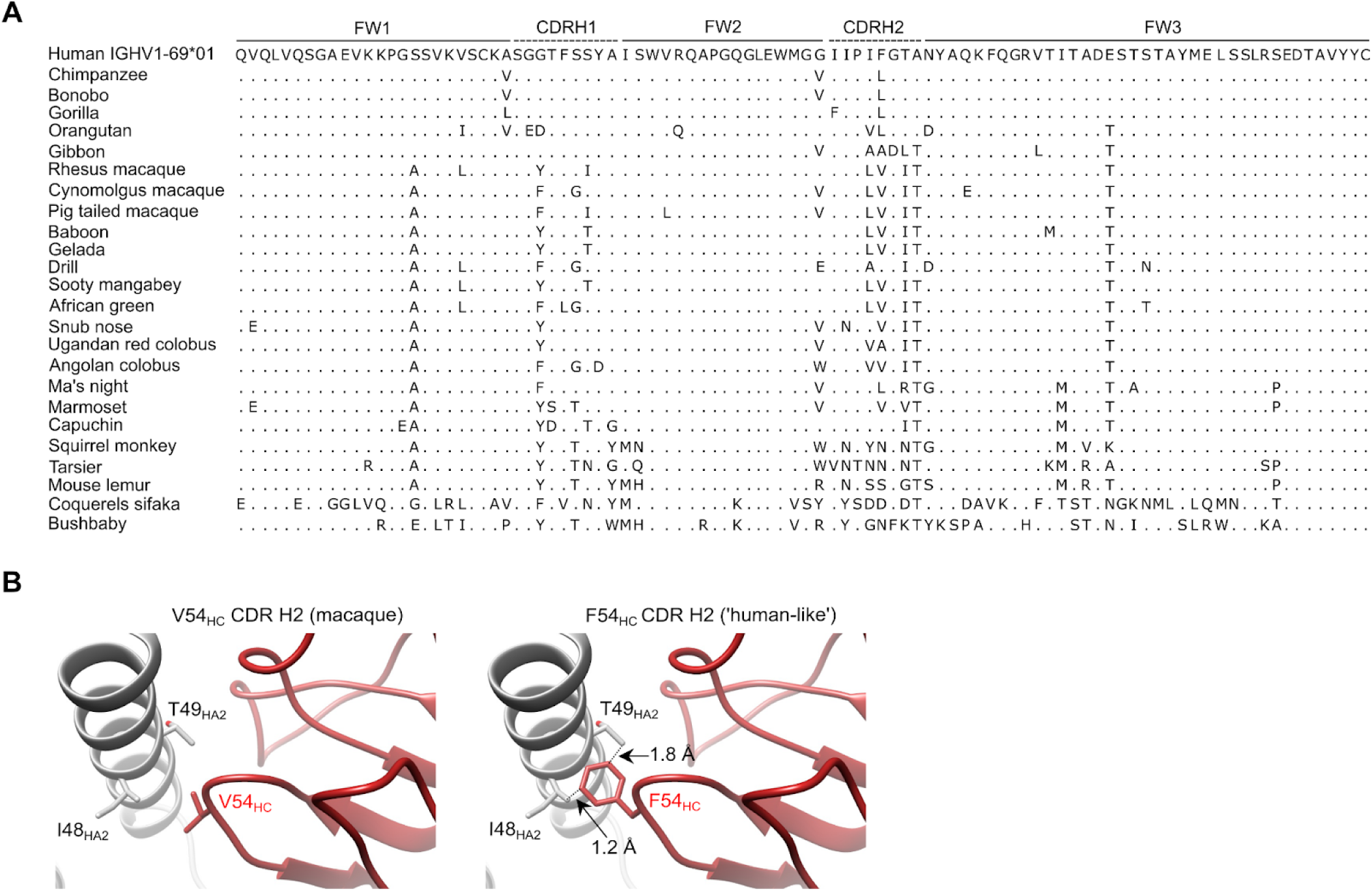
IGHV1-138 V_H_- gene characteristics across the primate order, related to Figure 5. **(A)** Amino acid alignment of germline human IGHV1-69*01V_H_-gene against non-human primate homologs. **(B)** Illustration of BB798E-3C07 interaction with HA highlighting the macaque CDR H2 V54_HC_ residue (Left), and a model showing insertion of CDR H2 F54_HC_ and steric clash to I48_HA2_ and I49_HA2_ (Right).

**Figure S6.**
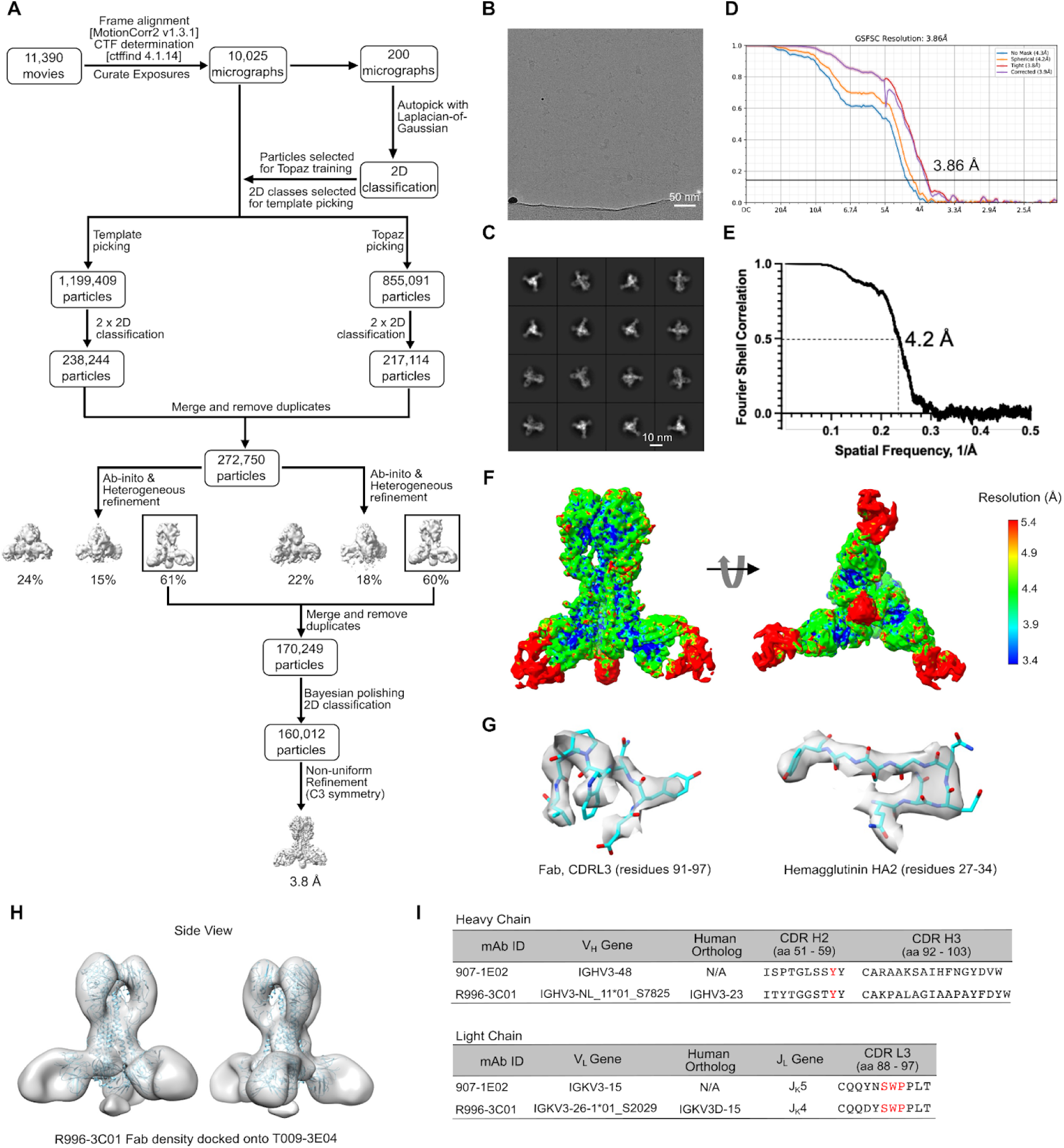
T009-3E04 Cryo-EM data processing and workflow, related to Figure 6. **(A)** Cryo-EM data processing and workflow for HA NC99 in complex with Fab T009 3E04. **(B-C)** Representative micrographs of 3E04 (B) and representative high-resolution 2D class averages of 3E04 (C). **(D)** Gold-standard resolution data generated by cryoSPARC. At the 0.143 threshold, the resolution is 3.86 Å for 3E04. **(E)** Fourier shell correlation curve between the map and the atomic model for 3E04. **(F)** Results of local resolution analysis using ResMap. The cryo-EM map is colored according to local resolution 3E04. **(G)** Examples of cryo-EM density for 3E04. **(H)** nsEM 3D reconstruction of nsEM R996-3C01 (SWP) Fab docked onto the cryo-EM structure of T009-3E04 (NWP) **(I)** HC and LC sequence characteristics of CDR L3 SWP mAbs from human (907-1E02) and macaque (R996-3C01). Sequence characteristics for macaque R996-3C01 are also shown in Figure 5A.

**Figure S7.**
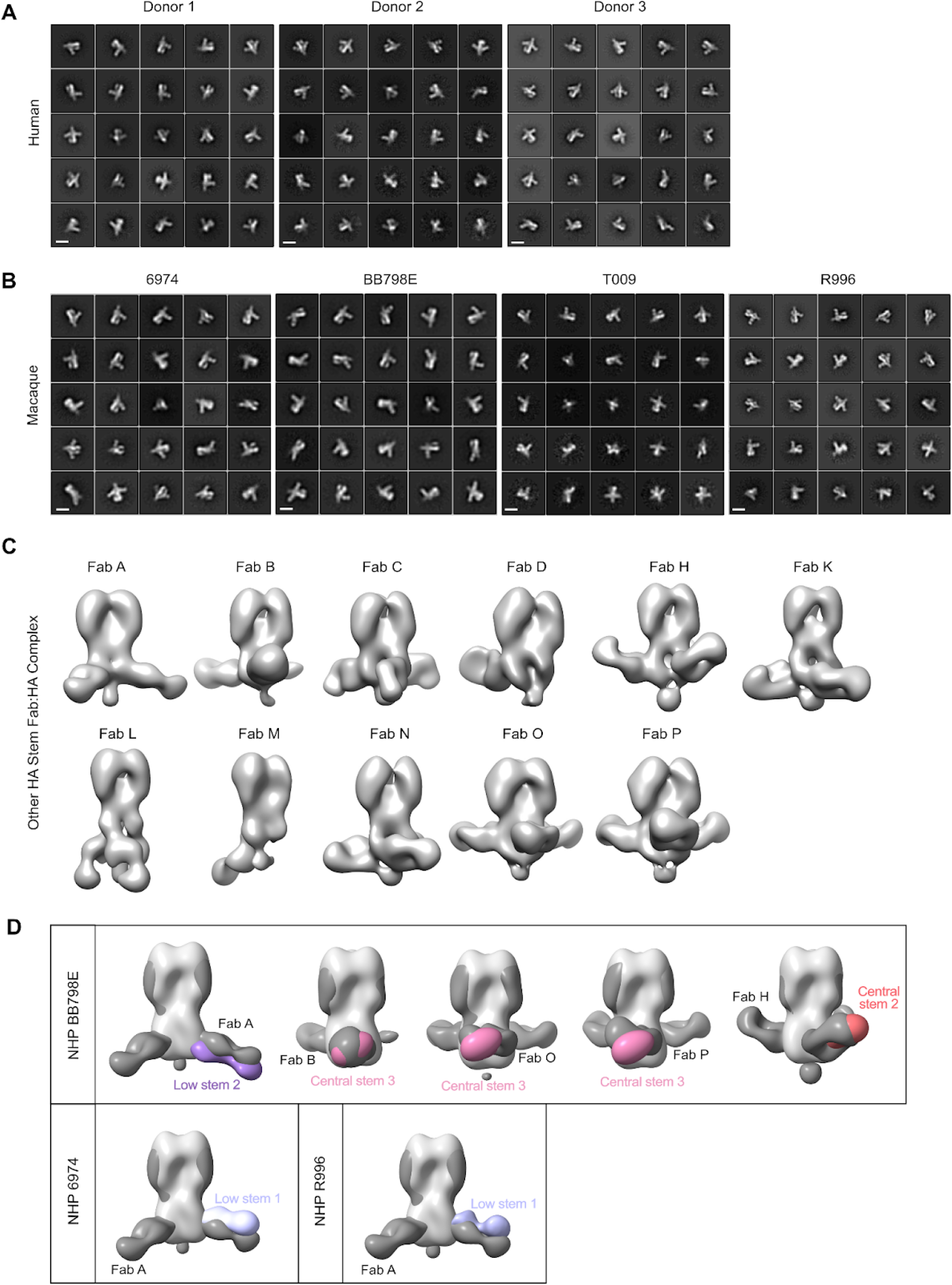
nsEMPEM polyclonal serum characterization, related to Figure 7. **(A-B)** nsEMPEM 2D class averages for each human (A) or macaques (B) serum antibody response. All 2D classification datasets are shown in order of descending particle count. Scale bar is 100Å. **(C)** nsEM and 3D reconstruction of other HA-stem macaque Fabs (see Figure 2F) in complex with H1 NC99 HA. **(D)** Overlay of other HA-stem macaque Fabs with relevant macaque nsEMPEM density (see Figure 7 C-D).

**Table S1.** Cryo-EM data collection and refinement statistics, Related to Figure 4 and 6.

|  | HA NC99/<br>Fab BB79E 3-C07 | HA NC99/<br>Fab T009 3-E04 |
| --- | --- | --- |
| **Data collection and processing** |  |  |
| Magnification | 22,500 | 22,500 |
| Voltage (kV) | 300 | 300 |
| Electron exposure (e<sup>-</sup>/Å<sup>2</sup>) | 40 | 40 |
| Defocus range (μm) | -0.9 to -2.3 | -1.0 to -2.7 |
| Pixel size (Å) | 1.11 | 1.11 |
| Symmetry imposed | C3 | C3 |
| Initial particle images (no.) | 371,568 | 2,054,500 |
| Final particle images (no.) | 104,541 | 160,012 |
|  | EMD-45636 | EMD-45637 |
|  | PDB 9CJY | PDB 9CJZ |
| Map resolution (Å) | 3.73 | 3.86 |
| FSC threshold | 0.143 | 0.143 |
| Map resolution range (Å) | 3.4-5.4 | 3.4-5.4 |
| **Refinement** |  |  |
| Initial model used (PDB code) | 8D21 | 8D21 |
| Model resolution (Å) | 4.1 | 4.2 |
| FSC threshold | 0.5 | 0.5 |
| Map sharpening *B* factor (Å<sup>2</sup>) | -140.5 | -158.8 |
| Model composition |  |  |
| Non-hydrogen atoms | 16318 | 15438 |
| Protein residues | 2050 | 1957 |
| Ligands | 0 | 0 |
| Water | 0 | 0 |
| *B* factors (Å<sup>2</sup>)(mean) |  |  |
| Protein | 41.98 | 168.01 |
| Ligand | N/A | N/A |
| Water | N/A | N/A |
| R.m.s. deviations |  |  |
| Bond lengths (Å) | 0.005 | 0.003 |
| Bond angles (°) | 0.597 | 0.510 |
| Validation |  |  |
| MolProbity score | 1.66 | 2.01 |
| Clash score | 5.34 | 10.59 |
| Poor rotamers (%) | 0.73 | 0.29 |
| Ramachandran plot |  |  |
| Favored (%) | 94.49 | 92.54 |
| Allowed (%) | 5.51 | 7.46 |
| Disallowed (%) | 0 | 0 |

